# Viscoelastic Extracellular Matrix Enhances Epigenetic Remodeling and Cellular Plasticity

**DOI:** 10.1101/2024.04.14.589442

**Authors:** Yifan Wu, Yang Song, Jennifer Soto, Tyler Hoffman, Xiao Lin, Aaron Zhang, Siyu Chen, Ramzi N. Massad, Xiao Han, Dongping Qi, Kun-Wei Yeh, Zhiwei Fang, Joon Eoh, Luo Gu, Amy C. Rowat, Zhen Gu, Song Li

**Affiliations:** Department of Bioengineering, University of California, Los Angeles, CA 90095, USA; Institute of Biomedical Engineering, West China School of Basic Medical Sciences & Forensic Medicine, Sichuan University, Chengdu 610041, China; Department of Integrative Biology and Physiology, University of California, Los Angeles, CA 90095, USA; Department of Materials Science and Engineering, Institute for NanoBioTechnology, and Translational Tissue Engineering Center, Johns Hopkins University, Baltimore, MD 21218, USA; California NanoSystems Institute, University of California Los Angeles, Los Angeles, CA 90095, USA; Jonsson Comprehensive Cancer Center, David Geffen School of Medicine, University of California Los Angeles, Los Angeles, CA 90095, USA; National Key Laboratory of Advanced Drug Delivery and Release Systems, College of Pharmaceutical Sciences, Zhejiang University, Hangzhou 310058, China; Key Laboratory of Advanced Drug Delivery Systems of Zhejiang Province, College of Pharmaceutical Sciences, Zhejiang University, Hangzhou 310058, China; MOE Key Laboratory of Macromolecular Synthesis and Functionalization, Department of Polymer Science and Engineering, Zhejiang University, Hangzhou 310027, China; Department of Medicine, University of California Los Angeles, Los Angeles, CA 90095, USA; Eli and Edythe Broad Center of Regenerative Medicine and Stem Cell Research, University of California Los Angeles, Los Angeles, CA 90095, USA

**Keywords:** matrix viscoelasticity, epigenetic state, chromatin dynamics, cell reprogramming

## Abstract

Extracellular matrices of living tissues exhibit viscoelastic properties, yet how these properties regulate chromatin and the epigenome remains unclear. Here, we show that viscoelastic substrates induce changes in nuclear architecture and epigenome, with more pronounced effects on softer surfaces. Fibroblasts on viscoelastic substrates display larger nuclei, lower chromatin compaction, and differential expression of distinct sets of genes related to the cytoskeleton and nuclear function compared to those on purely elastic surfaces. Slow-relaxing viscoelastic substrates reduce lamin A/C expression and enhance nuclear remodeling. These structural changes are accompanied by a global increase in euchromatin marks and local increase in chromatin accessibility at cis-regulatory elements associated with neuronal and pluripotent genes. Consequently, viscoelastic substrates improve the reprogramming efficiency from fibroblasts into neurons and induced pluripotent stem cells. Collectively, our findings unravel the roles of matrix viscoelasticity in epigenetic regulation and cell reprogramming, with implications for designing smart materials for cell fate engineering.

## Main

Native tissue and the extracellular matrix (ECM) are biomolecular assemblies possessing viscoelastic properties^1^. With the development of biomaterials with tunable mechanical properties^2,3^, viscoelastic effects have been shown to regulate both cell-level processes such as spreading, migration, and differentiation^4–6^, and tissue-level behaviors such as matrix formation, morphogenesis, and organoid development^7–10^, which underscores the important roles of matrix viscoelasticity in development, disease progression, and tissue regeneration. Although several studies have investigated the cytoskeletal reorganization on viscoelastic substrates to connect extracellular mechanical changes with intracellular signaling, the roles of the cell nucleus as an important mechanosensory unit in response to matrix viscoelasticity is largely unknown.

Accumulative evidence has shown that mechanical stimuli, such as compression and matrix stiffness, can regulate chromatin and the epigenome in the cell nucleus to influence the on-off state of lineage-specific genes and thus, cell fates^11–17^. Deciphering whether and how matrix viscoelasticity affects the cell nucleus to modulate cell fate, especially chromatin and the epigenome, will shed lights on the development of smart materials to control cell phenotype conversion. To achieve this, we engineered the stiffness and viscoelasticity of cell-adhesive substrates, and investigated the genome-wide biophysical and biochemical changes globally and at specific sites. Our findings provide a mechanistic interpretation of how the cell nucleus responds to matrix viscoelasticity to regulate chromatin and epigenome and thus, modulate cellular plasticity, potentially leading to a new technology platform to engineer cell fate for tissue regeneration, disease modeling, and drug screening.

### Stiffness-dependent viscoelastic effects on cell proliferation and spreading

Soft biological tissues, such as brain, abdominal organs, and skin, have elastic moduli on the kilopascal level and stress relaxation half times less than thousands of seconds^1,18^. To mimic their mechanical properties, we fabricated three groups of alginate-based hydrogels with initial elastic moduli of 2, 10, and 20 kPa and stress relaxation half times (*τ*_½_, the time under constant deformation to relax the initial stress to half of the original value) of ∼200 and ∼1000 s as described previously^6^. As alginate gels allow users to tailor viscoelasticity independent of stiffness, pore size, and ligand density^19,20^, this system can dissect the influence of matrix stiffness and viscoelasticity on cell behaviors.

By varying the type and concentration of crosslinkers and the molecular weight of alginate polymers, we obtained alginate gel substrates of 2, 10, and 20 kPa with little stress relaxation behavior (covalent crosslinking, elastic), and slow or fast relaxation (ionic crosslinking, slow relaxing or fast relaxing, viscoelastic), respectively (**Fig. 1a-c**). Characterization of the derived gels showed that while 2-kPa elastic and slow-relaxing gels swelled after 1 day of DMEM (Dulbecco’s Modified Eagle’s Medium) immersion compared to other gels, there was no significant volume changes between 1 and 7 days in all the gels examined (**Supplementary Fig. 1a**). Additionally, the dry mass of all gel substrates was stable for at least seven days under tissue culture (TC) conditions (**Supplementary Fig. 1b**), proving its suitability as a long-term cell culture platform. Rheology measurements showed that gels formed by covalent crosslinking had constant elastic modulus under different frequencies and near-zero loss modulus (**Fig. 1b, Supplementary Fig. 1c**). Conversely, the ionic crosslinking afforded gels frequency-dependent elastic moduli and significantly higher loss moduli, where fast-relaxing gels displayed higher loss moduli than slow-relaxing gels at most frequencies (**Fig. 1b, Supplementary Fig. 1c**), showing well-defined viscoelastic properties. Consistently, compression tests demonstrated that ionically crosslinked gel substrates (slow relaxing and fast relaxing) displayed different degrees of stress-relaxation under deformation (**Fig. 1c**, **Supplementary Fig. 1d**). Moreover, we confirmed that viscoelastic properties of the gels are independent of stiffness, as stress relaxation half time was similar across different stiffnesses for both slow-relaxing (*τ*_½_ ∼1000 s, **Supplementary Fig. 1e**) and fast-relaxing gels (*τ*_½_ ∼200 s, **Supplementary Fig. 1e**).

**Figure 1.**
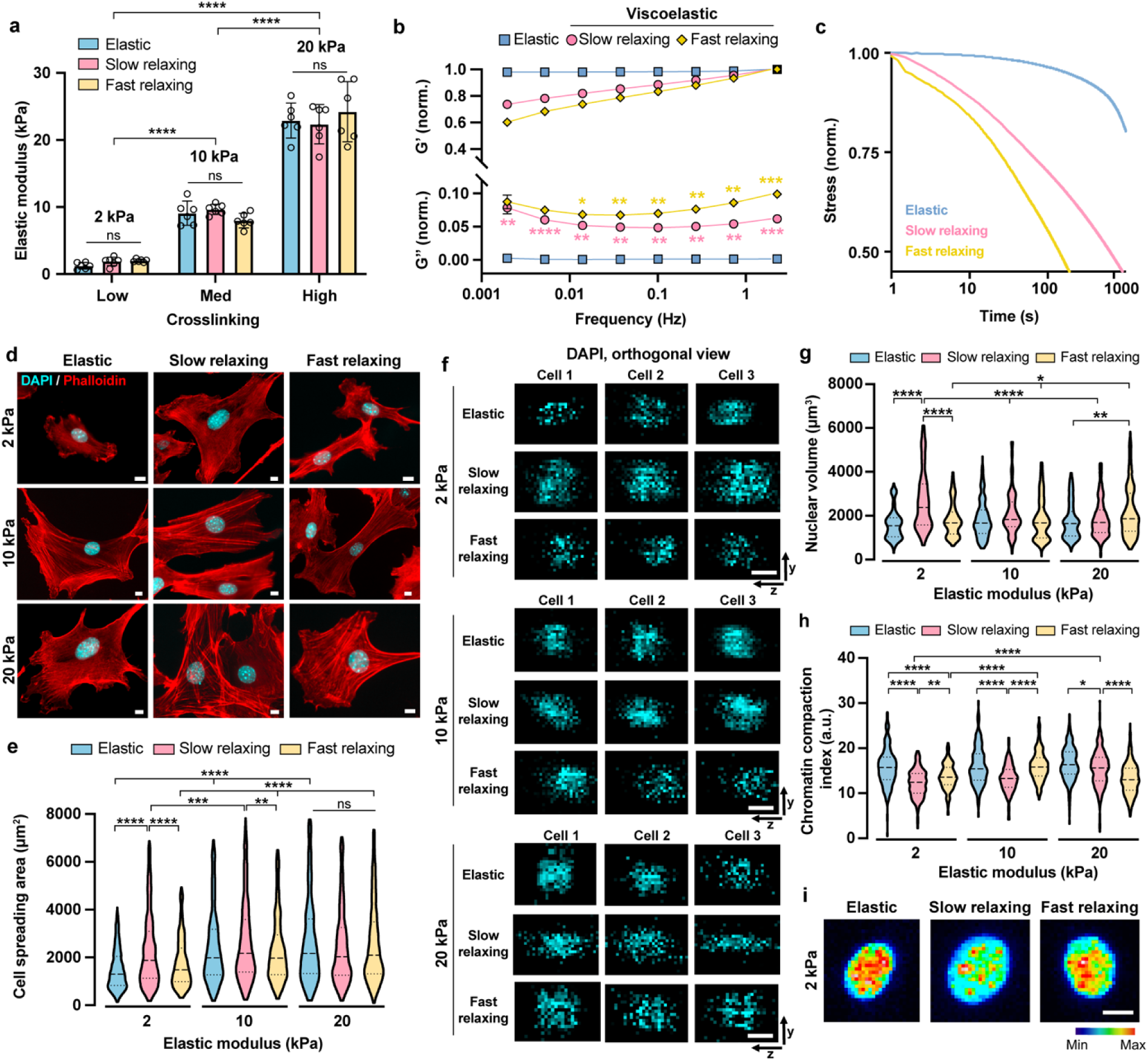
Viscoelastic substrates regulate cell spreading, nuclear volume, and chromatin compaction in a stiffness-dependent manner. **a**, Initial elastic modulus of covalently (elastic) or ionically (viscoelastic, slow relaxing and fast relaxing) crosslinked alginate hydrogels with different concentrations of crosslinker, as measured by compression tests (n = 6 batches of gels). **b**, Storage modulus (G’) and loss modulus (G’’) of alginate gels at varying frequencies (n = 3 batches of gels). Values are normalized by the storage modulus at 1 rad s^−1^ of the gels presented. **c**, Stress relaxation of gels as a function of time under a constant deformation. **d**, Representative fluorescent images of the cytoskeletal network in mouse fibroblasts on different substrates after 24 hours, where F-actin (phallodin, red) and nuclei (DAPI, blue) are labeled. Scale bar, 10 *μ*m. **e**, Cell spreading area on various substrates (n= 810, 676, 381, 608, 893, 489, 589, 831, 604 cells per condition from 3 independent experiments). **f**, Orthogonal view of three representative cell nuclei on various substrates. Scale bar, 10 *μ*m. **g**, Nuclear volume of fibroblasts cultured on gel substrates for 24 hours (n = 187, 140, 167, 154, 136, 173, 193, 346, 174 cells per condition from 3 independent experiments). **h**, Chromatin compaction index of fibroblasts on gels at 24 hours, quantified by the ratio of integrated fluorescence intensity of DAPI staining to nuclear volume (n = 186, 140, 163, 152, 133, 172, 193, 344, 174 cells per condition from 3 independent experiments). **i**, Representative images of a DAPI-stained cell nucleus on 2-kPa substrates. Scale bar, 10 *μ*m. In **a** and **b**, data are shown as mean ± s.d. In **e**, **g**, and **h**, truncated violin plots were used to demonstrate the distribution of data. The violins are drawn with the ends at the quartiles and the median as a horizontal line in the violin, where the curve of the violin extends to the minimum and maximum values in the data set. Significance was determined by one-way ANOVA using Tukey’s correction for multiple comparisons (\**P* < 0.05, \*\**P*<0.01, \*\*\**P* <0.001, \*\*\*\**P* < 0.0001).

Fibroblasts are the most abundant cell type in connective tissue throughout the body. It plays important roles in tissue homeostasis and serve as major cell sources for direct cell reprogramming to generate functional cells with therapeutic potentials^21,22^. As native tissue and ECMs are viscoelastic, understanding whether the cellular plasticity of fibroblasts is regulated by matrix viscoelasticity is of great promise to advance our knowledge of tissue biology and regenerative medicine. Primary fibroblasts isolated from adult mice were seeded on the alginate-based hydrogels made from RGD (Arg-Gly-Asp)-coupled alginate polymers with covalent or ionic crosslinking, where RGD peptides served as ligands for cell adhesion and spreading^20^. We found that cell viability on the gels was nearly 100% as on TC plates at 24, 48, and 72 hours after cell seeding (**Supplementary Fig. 2a-d**). Cell proliferation analysis utilizing 5-ethynyl-2’-deoxyuridine (EdU) labeling at 48 hours after cell seeding showed that on purely elastic surfaces, cell proliferation increased with stiffness (**Supplementary Fig. 2e-f**). Viscoelastic surfaces increased cell proliferation on 2-kPa and 10-kPa substrates, compared to purely elastic surfaces. In contrast, no significant difference was seen on 20-kPa substrates regardless of viscoelasticity (**Supplementary Fig. 2e-f**), indicating that matrix viscoelasticity promotes cell proliferation on softer surfaces (2 kPa and 10 kPa), but not stiff surfaces (20 kPa).

We then investigated how viscoelasticity regulated cell spreading on substrates with different stiffness. Fibroblasts could not fully spread on soft elastic surfaces (2 kPa) as was evident on stiffer elastic surfaces of 10 kPa and 20 kPa (**Fig. 1d-e**). Interestingly, cells spread more on 2-kPa slow-relaxing gels, but not on fast-relaxing gels, compared to those on 2-kPa elastic gels (**Fig. 1d-e**, **Supplementary Fig. 3a**). A possible explanation is that cell adhesion formation on soft substrates is associated with time-dependent force responses^11,23^. On slow-relaxing substrates of 2 kPa (*τ*_½_ ∼1000 s), a low level of traction force affords sufficient time for the cells to form focal adhesions, while cells on 2-kPa substrates with faster relaxation (*τ*_½_ ∼200 s) may experience a higher ratio of focal adhesion turnover to formation. In contrast, on 10-kPa and 20-kPa substrates, viscoelasticity did not further facilitate cell spreading (**Fig. 1d-e**, **Supplementary Fig. 3a**).

### Stiffness-dependent viscoelastic effects on the cell nucleus

The cell nucleus is an important mechanosensor and processor ^24–28^. To determine whether viscoelastic substrates impact cell nucleus, we quantified the nuclear volume of cells cultured on the various substrates. Different from cell proliferation and spreading, nuclear volume was not significantly affected by stiffness on elastic surfaces, but was sensitive to substrate viscoelasticity (**Fig. 1f-g**, **Supplementary Figs. 3b and 4**). Cells cultured on slow-relaxing gels showed an increase in nuclear volume only on soft (2 kPa) surfaces compared to purely elastic substrates, while cells on fast-relaxing gels had increased nuclear volume on stiff surfaces (20 kPa) but not on 2-kPa and 10-kPa substrates. These findings suggest that matrix viscoelasticity modulates the physical properties of the cell nucleus, which is dependent on stiffness and relaxation time.

We then directly investigated whether matrix viscoelasticity regulates chromatin compaction by examining the degree of chromatin condensation inside cell nuclei on various substrates. By calculating the ratio of integrated 4′,6-diamidino-2-phenylindole (DAPI) fluorescence intensity to nuclear volume in each cell as described previously^29,30^, we obtained the chromatin compaction index, where lower value indicates less chromatin condensation. We found that fibroblasts on slow-relaxing gels displayed lower chromatin compaction compared to those on elastic gels across different stiffnesses, where the most pronounced differences were observed on 2-kPa slow-relaxing substrates (**Fig. 1h-i**, **Supplementary Figs. 3c and 5**). Additionally, cells on 20-kPa fast-relaxing gels had much lower chromatin compaction indices than those on elastic and slow-relaxing gels, (**Fig. 1h-i**, **Supplementary Figs. 3c and 5**).

### Substrate viscoelasticity induces distinct gene expression profiles compared with substrate stiffness

It has been shown that chromatin architecture impacts gene expression^31^. As chromatin was less condensed in fibroblasts on viscoelastic substrates, we hypothesized that there were accompanying transcriptomic changes in response to substrate viscoelasticity. Therefore, we performed RNA-sequencing (RNA-seq) on fibroblasts seeded on various substrates at 48 hours to determine whether substrate viscoelasticity altered gene expression in a stiffness-dependent manner. Considering the pronounced effects of slow-relaxing substrates on cell proliferation, spreading, nuclear volume, and chromatin compaction, together with the stiffness dependence of these viscoelasticity-induced changes, we employed elastic and slow-relaxing gels of 2 kPa and 20 kPa as representatives for RNA-seq analysis.

Principal component analysis (PCA) results demonstrated that fibroblasts growing on elastic and viscoelastic substrates were clustered at different sides of the principal component 1 (PC1) axis, which suggests that viscoelastic substrates regulate different sets of genes from elastic substrates (**Fig. 2a**). Interestingly, while cells on 2-kPa and 20-kPa elastic gels can be found at different ends of the PC2 axis illustrating stiffness effects, cells on viscoelastic substrates of these two stiffnesses were close along the PC2 axis. This proximity was confirmed by the substantially fewer differentially expressed genes (DEGs) between cells on 2-kPa and 20-kPa slow-relaxing gels, compared to other groups (**Fig. 2b**). These findings suggest that while viscoelastic substrates influence gene expression differently compared to stiffness alone, viscoelasticity results in similar transcriptomic profiles in cells on both soft and stiff surfaces. This indicates that substrate stiffness has a reduced impact in the presence of viscoelasticity.

**Figure 2.**
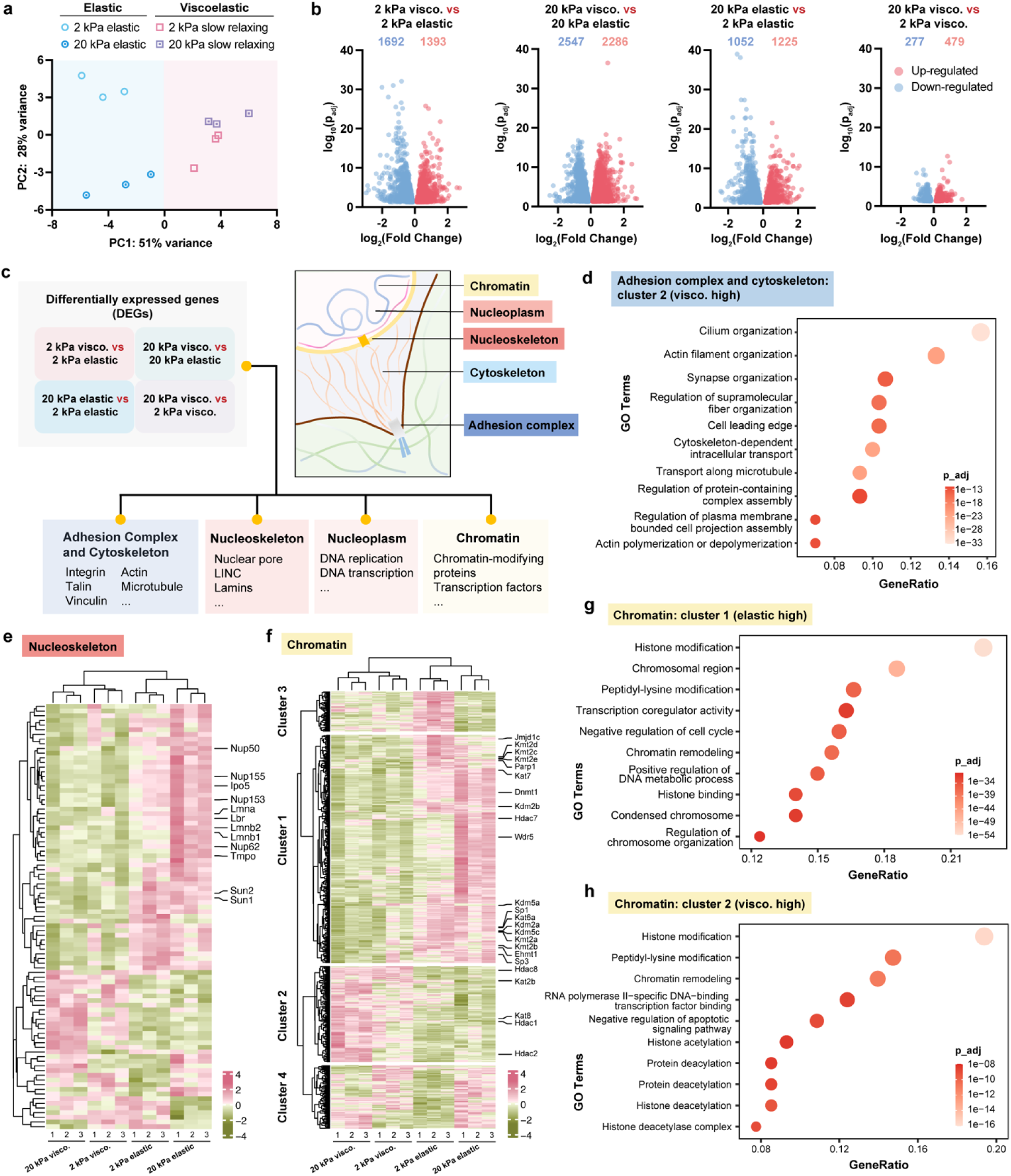
RNA-seq of fibroblasts on gel substrates with varying stiffness and viscoelasticity. **a**, Principal component analysis (PCA) of RNA-seq data of mouse fibroblasts cultured on different gels for 48 hours (n = 3 biological replicates). **b**, Volcano plots of differentially expressed genes (DEGs). The number of up- and down-regulated genes in each group are indicated using red and blue numbers. **c**, Schematic representation of DEGs from the categorized mechanotransduction-related gene database. **d**, Gene ontology (GO) enrichment analysis of genes up-regulated in viscoelastic groups in the Adhesion complex and cytoskeleton category. **e-f**, Heatmaps of DEGs in the Nucleoskeleton (**e**) and Chromatin (**f**) categories. **g-h**, GO terms enriched in genes up-regulated in elastic (**g**) and viscoelastic (**h**) groups from the Chromatin category.

To determine viscoelasticity-induced transcriptomic changes in mechanotransduction, we established a mechanotransduction-related gene regulatory network database and classified relevant genes into 4 categories (adhesion complex and cytoskeleton, nucleoskeleton, nucleoplasm, and chromatin) (**Fig. 2c**). By generating DEG expression heatmaps by categories, we found that these category-specific genes can be grouped into 3 or 4 clusters containing genes up-regulated on elastic groups (cluster 1), genes up-regulated on viscoelastic groups (cluster 2), and genes up- or down-regulated only on 2-kPa elastic gels (cluster 3 and cluster 4, respectively). We then performed gene ontology (GO) enrichment analysis to provide functional annotations for these clusters.

In the ‘Adhesion complex and cytoskeleton’ category, integrin-related genes and genes encoding vinculin and talin were down-regulated in cells on viscoelastic surfaces (**Supplementary Fig. 6a**), suggesting that there may be alterations in cell adhesion strength on substrates without and with stress relaxation behaviors. GO terms related to cytoskeletal filament organization and intracellular transport were enriched in genes up-regulated in viscoelastic groups (**Fig. 2d**). Additionally, several microtubule-related genes were up-regulated in the viscoelastic groups (**Supplementary Fig. 6a**), which implied the potential role of microtubules in transducing substrate viscoelasticity, alongside the previously reported actin-related pathways^6^.

The gene expression patterns in the Nucleoskeleton, Nucleoplasm, and Chromatin categories demonstrated that viscoelastic surfaces had distinct effects on the cell nucleus compared to purely elastic surfaces (**Fig. 2e-f**, **Supplementary Fig. 6b**). For example, cells cultured on soft elastic gels (2 kPa) showed reduced expression of genes encoding lamins, linker of nucleoskeleton and cytoskeleton (LINC) complex proteins, and nuclear pore complex proteins, compared to stiff elastic gels (20 kPa) (**Fig. 2e**). The expression of these genes was further decreased on viscoelastic surfaces (**Fig. 2e**). In addition, the levels of mRNAs that encode genes related to epigenetic remodeling, such as various chromatin-modifying enzymes, changed among elastic and viscoelastic groups (**Fig. 2f**). Elastic surfaces promoted genes related to condensed chromosome as shown in the GO analysis plot (**Fig. 2g**), which was consistent with our findings in chromatin condensation. It is also worth noting that viscoelastic surfaces enhanced the expression of genes involved in histone acetylation (**Fig. 2h**). Taken together, these findings provide insights into the possible mechanisms of substrate viscoelasticity-mediated epigenetic remodeling.

### Viscoelastic substrates promote cell reprogramming from fibroblasts to induced neurons and induced pluripotent stem cells (iPSCs)

Considering that viscoelastic substrates regulate various aspects of the cell nucleus related to the epigenetic state, such as chromatin condensation and genes involved in nuclear lamina structure and histone acetylation, we postulated that a viscoelastic substrate would modulate cellular plasticity and thus, impact the cell reprogramming process. Therefore, we employed a direct cell reprogramming model that converted mouse fibroblasts to neurons by lentiviral transduction of three transcription factors (Brn2, Ascl1, Myt1l, abbreviated as BAM)^32^. To test the effect of substrate viscoelasticity on induced neuronal (iN) reprogramming, primary mouse fibroblasts were transduced with doxycycline (Dox)-inducible lentiviral vectors encoding BAM, and the following day, the cells were seeded onto elastic and viscoelastic substrates with the same initial cell density. Twenty-four hours later, Dox was added to the culture media to induce the expression of the transgenes and initiate the reprogramming process (marked as day 0). Neuronal medium containing Dox was used for cell culture from day 1 to the conclusion of the experiment (**Fig. 3a**).

**Figure 3.**
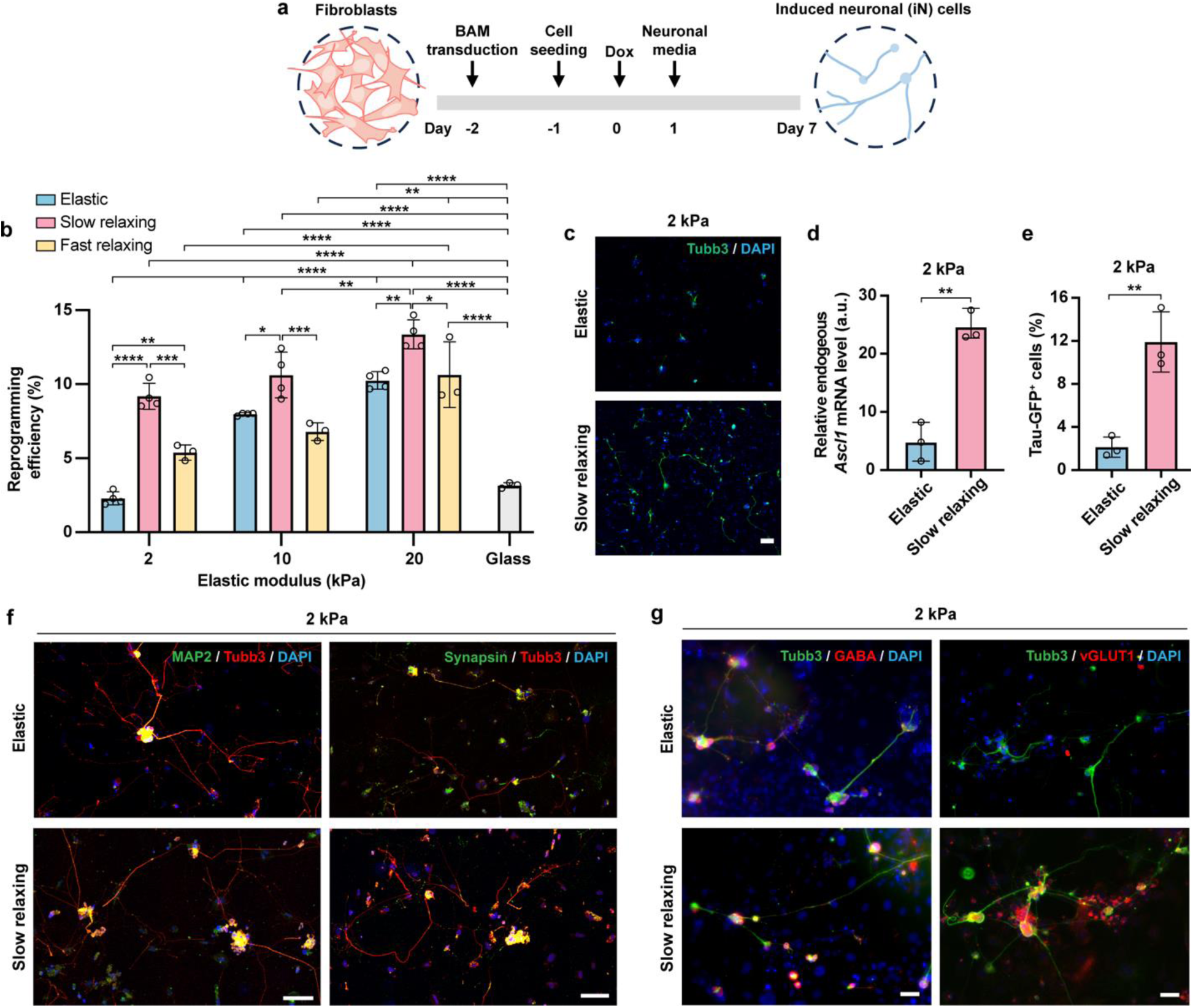
Substrate viscoelasticity enhances the reprogramming of fibroblasts into induced neuronal (iN) cells. **a**, Schematic illustration of the reprogramming timeline. **b**, Reprogramming efficiency of BAM-transduced fibroblasts cultured on various substrates at day 7, calculated as the percentage of Tubb3^+^ cells among the total number of initially seeded cells (n = 4, 4, 3, 3 independent experiments for elastic, slow-relaxing, fast-relaxing gels, and glass). **c**, Representative fluorescent images showing Tubb3^+^ cells at day 7 on 2-kPa substrates. Scale bar, 100 *μ* m. **d**, Endogenous *Ascl1* mRNA expression in BAM-transduced cells on 2-kPa substrates at day 2 after doxycycline (Dox)-induced transgene expression (n = 3 independent experiments). **e**, Percentage of cells expressing Tau-GFP on different substrates at day 5 (n = 3 independent experiments). **f**, Representative fluorescent images showing Tubb3^+^ iN cells co-expressing mature neuronal markers, MAP2 or synapsin at day 21. Scale bar, 100 *μ*m. **g**, Representative fluorescent images showing Tubb3^+^ iN cells co-expressing gamma-aminobutyric acid (GABA) or vesicular glutamate transporter 1 (vGLUT1) at day 28. Scale bar, 50 *μ*m. In **b** and **d-e**, bar graphs show the mean ± s.d. In **b**, a two-way ANOVA using Tukey’s correction for multiple comparisons was used to determine the significance between the groups. In **d-e**, a two-tailed, unpaired t test was used to determine statistical significance (\**P* < 0.05, \*\**P* < 0.01, \*\*\**P*<0.001, \*\*\*\**P* < 0.0001).

To determine the reprogramming efficiency of iN conversion, we fixed the samples on day 7, immunostained the cells for neuronal marker Tubb3 (neuron-specific class III beta-tubulin), and calculated the percentage of Tubb3^+^ cells among the total number of initially seeded cells. On elastic surfaces, reprogramming efficiency increased with the stiffness, in agreement with previous findings^16^ (**Fig. 3b**). Interestingly, slow-relaxing substrates led to a significantly higher iN conversion efficiency compared to elastic substrates with the same stiffness (**Fig. 3b-c**), with the most pronounced effects on 2-kPa surfaces (∼4-fold increase) in comparison with stiffer surfaces (10 and 20 kPa, ∼30 % increase) (**Fig. 3b**). Compared with elastic surfaces, fast-relaxing gels only enhanced the reprogramming efficiency on 2-kPa surfaces (∼2 fold) (**Fig. 3b**). Therefore, we used 2-kPa elastic and slow-relaxing gels as representative substrates for further investigation of the reprogramming mechanisms.

Since Ascl1 serves as a pioneer factor in iN reprogramming that immediately binds to its genomic sites after Dox induction and recruits other transcription factors^33^, achieving a high level of endogenous *Ascl1* gene activation is critical for efficient iN reprogramming. On day 2, slow-relaxing gels induced a ∼5-fold increase of endogenous *Ascl1* expression in fibroblasts compared to elastic substrates, as measured by quantitative polymerase chain reaction (qPCR) using primers that could distinguish endogenous and transduced *Ascl1* gene (**Fig. 3d**). This finding indicates that slow-relaxing gels facilitate the activation and expression of neuronal genes in the heterochromatin of fibroblasts. Furthermore, we isolated fibroblasts from transgenic mice that expressed enhanced green fluorescence protein (EGFP) driven by the promoter of neuron-specific microtubule-associated protein (Tau), and conducted iN reprogramming experiments following the same procedure. On day 5, the percentage of cells expressing Tau-GFP on slow-relaxing substrates was ∼6-fold higher than those on elastic substrates (**Fig. 3e**), further confirming that slow-relaxing substrates promote iN reprogramming with higher efficiency.

To confirm the functionality of iN cells obtained from these substrates, we cultured iN cells for 3-4 weeks and found that iN cells expressed mature neuronal markers, microtubule-associated protein 2 (MAP2) and synapsin, on both surfaces after 3 weeks (**Fig. 3f**). After 4 weeks, they expressed gamma-aminobutyric acid (GABA) and vesicular glutamate transporter 1 (vGLUT1) on both substrates (**Fig. 3g**), indicative of the GABAergic and glutamatergic subtypes, consistent with previous reports^32,34^. Additionally, iN cells reprogrammed from fibroblasts on these substrates exhibited rhythmic calcium oscillations (**Supplementary Videos 1-2**), a characteristic of functional mature neurons, and similar morphology to neuronal cells derived on glass (**Supplementary Fig. 7**). Based on these findings, we concluded that viscoelastic gels boosted the reprogramming of adult mouse fibroblasts into matured neurons.

To further verify the effects of viscoelastic substrates on cell reprogramming, an additional reprogramming model was utilized by converting the fibroblasts into iPSCs. We transduced primary adult mouse fibroblasts with Oct4, Sox2, Klf4, and c-MYC (OSKM) transcription factors and cultured them on gel substrates with varying stiffness and viscoelasticity for 10 days (**Supplementary Fig. 8a**). Similarly, on elastic substrates, reprogramming efficiency increased with stiffness between 2 and 20 kPa^16^ (**Supplementary Fig. 8b**), as determined by the number of Nanog^+^ colonies. In addition, both slow-relaxing and fast-relaxing gels increased the iPSC reprogramming efficiency by ∼5-fold in comparison with elastic gels, which was only evident on 2-kPa substrates but not on 10-kPa and 20-kPa substrates (**Supplementary Fig. 8b**). These results show the same trends as iN reprogramming, and underscore the more pronounced effects of viscoelasticity on soft (2 kPa) substrates. In addition, Nanog^+^ colonies were larger in size on slow-relaxing substrates than elastic substrates of 2 kPa (**Supplementary Fig. 8c**), implying an accelerated iPSC reprogramming process. These findings were further supported by PCR analysis results that demonstrated a significant increase in *Nanog* expression from cells cultured on slow-relaxing substrates compared to elastic substrates of 2 kPa (**Supplementary Fig. 8d**). To verify the functionality of iPSCs generated on these substrates, we isolated the iPSCs from 2-kPa elastic and slow-relaxing substrates, and confirmed that they had the differentiation potential to develop into cells of all three germ layers, consistent with iPSCs generated on TC plates (**Supplementary Fig. 8e**).

### Substrate viscoelasticity facilitates nuclear lamina remodeling

Since the effects of substrate viscoelasticity had more pronounced effects on soft (2 kPa) substrates for both iN and iPSC reprogramming, we focused on 2-kPa gel substrates to further study the regulation of nuclear alternations and cell reprogramming by viscoelasticity. The nuclear lamina, which is mainly composed of A-type and B-type lamins, plays a major role in mechanosensing. From the RNA-seq analysis, we observed that cells on viscoelastic substrates exhibited down-regulated gene expression of *Lmna,* encoding A-type lamins, but there was no change in *Lmnb1,* which encodes one of the two B-type lamins, in fibroblasts on 2-kPa viscoelastic gels compared to purely elastic ones (**Fig. 2e**). Since the nuclear lamina is critical to convey the extracellular mechanical perturbations into the cell nucleus^35^, we investigated the effects of substrate viscoelasticity on protein expression, structure, and dynamics of lamins.

We first determined whether the protein expression of lamin A/C and lamin B1 followed the same patterns as the gene expression profile on substrates with different stress relaxation behaviors. Western blotting results showed that fibroblasts on 2-kPa slow-relaxing gels (but not fast relaxing gels) expressed a lower amount of lamin A/C compared to those on 2-kPa elastic gels, while lamin B1 expression remained similar on all 2-kPa substrates (**Fig. 4a-d**). In agreement with previous reports that lamin A/C-deficient fibroblasts exhibited reduced nuclear stiffness^36,37^, we also observed that cells on 2-kPa slow-relaxing substrates had softer nuclei than those on 2-kPa elastic gels (**Supplementary Fig. 9a**), suggesting that there may be an increase in the deformability of cell nuclei on slow-relaxing substrates.

**Figure 4.**
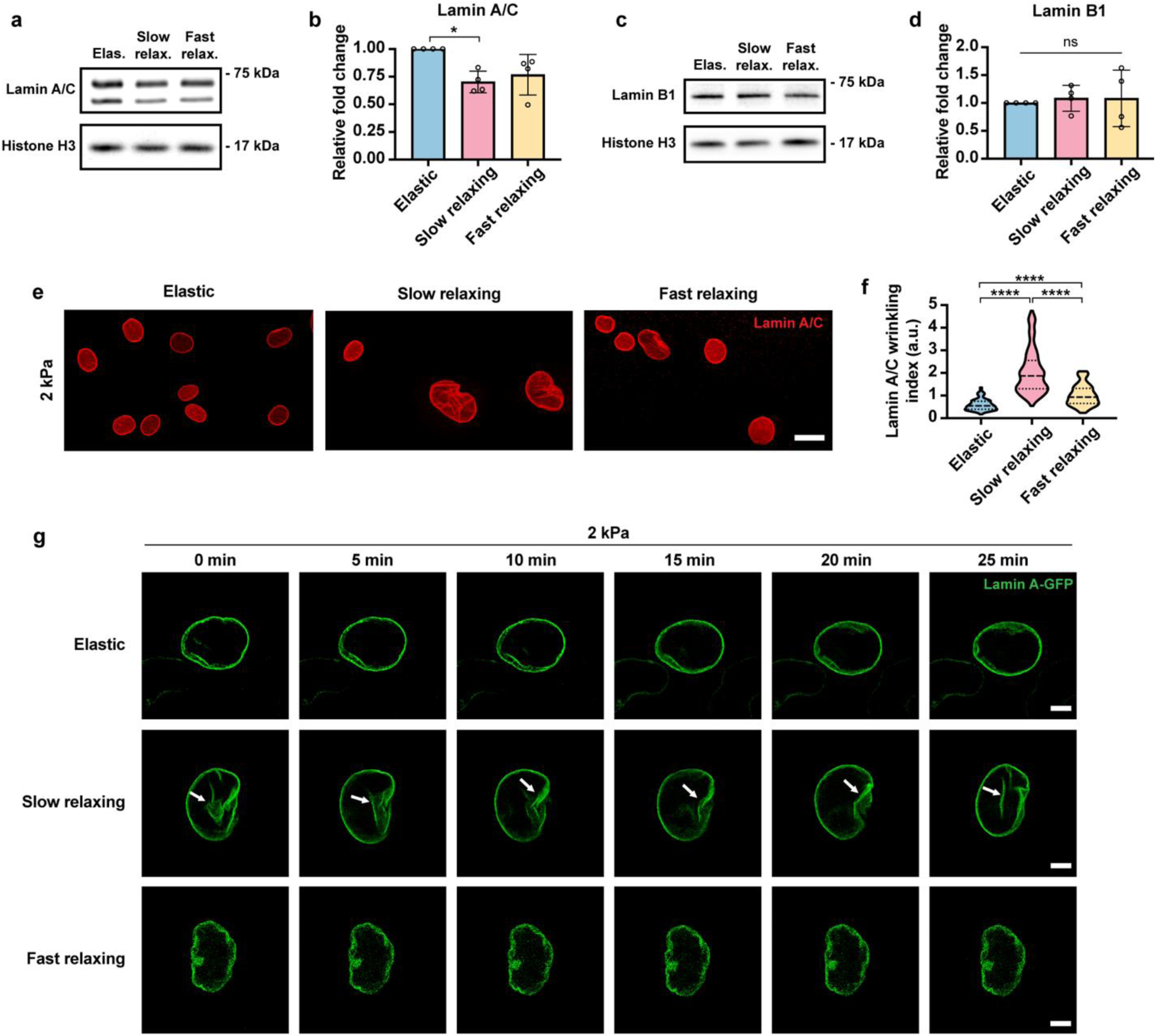
Viscoelastic substrates remodel the nuclear lamina. **a**, Western blot analysis of lamin A/C expression in fibroblasts cultured on various substrates for 48 hours, where histone H3 serves as the loading control. **b,** Quantification of lamin A/C expression from blots in **a**, where the protein expression level was normalized to the corresponding histone H3 and elastic groups (n = 3 independent experiments). **c**, Western blot analysis of lamin B1 expression in fibroblasts cultured on various substrates for 48 hours, where histone H3 serves as the loading control. **d,** Quantification of lamin B1 expression from blots in **c**, where the protein expression level was normalized to the corresponding histone H3 and elastic groups (n = 3 independent experiments). **e**, Immunostaining of lamin A/C in fibroblasts on 2-kPa substrates. Scale bar, 20 *μ*m. **f,** Quantification of nuclear wrinkling index based on lamin A/C staining (n = 146, 145, 116 cells per condition from 3 independent experiments). **g**, Time-lapse microscopy images of fibroblasts expressing GFP-tagged lamin A on 2-kPa substrates. Scale bar, 5 *μ*m. In **b** and **d**, data are shown as mean ± s.d. In **f**, the violins are drawn with the ends at the quartiles and the median as a horizontal line in the violin, where the curve of the violin extends to the minimum and maximum values in the data set. Significance was determined by one-way ANOVA using Tukey’s correction for multiple comparisons (\**P* < 0.05, \*\*\*\**P* < 0.0001).

Next, to investigate the structural changes of the nuclear lamina induced by substrate viscoelasticity, fibroblasts on various substrates were immunostained for lamin A/C and lamin B1 at 48 hours after cell seeding. As shown by lamin A/C staining, cells on 2-kPa viscoelastic gels displayed increased nuclear wrinkling compared to those on 2-kPa elastic substrates, where slow-relaxing substrates enhanced nuclear wrinkling more than fast-relaxing gels (**Fig. 4e-f**, **Supplementary Figs. 9b and 10**). Lamin B1 staining revealed similar nuclear wrinkling patterns (**Supplementary Figs. 9c and 10**). To be noted, lamin B1 and lamin A/C did not fully overlap in the nucleoplasm in fibroblasts on different substrates (**Supplementary Figs. 9d and 10**), leading to variation in defining nuclear wrinkling patterns based on different markers. These findings also highlight the heterogeneity of nuclear lamin networks and the distinct functions of A-type and B-type lamins in mechanotransduction^38^.

Moreover, to observe the spatiotemporal changes of the nuclear lamina in fibroblasts on gel substrates with varying viscoelasticity, we recorded the activity of fibroblasts expressing green fluorescence protein (GFP)-tagged lamin A on 2-kPa gels using time-lapse fluorescence microscopy (**Fig. 4g**, **Supplementary Video 3-8**). We found that there were more dynamic changes in the nuclear lamina (e.g., deformation and the appearance and disappearance of nuclear wrinkles) of cells on viscoelastic substrates, especially slow-relaxing gels, possibly due to the dynamic stress relaxation and resulting intracellular stress changes around the cell nucleus. Together, these findings suggest that a viscoelastic substrate may facilitate the dynamic remodeling of the nucleus.

### Viscoelastic substrates regulate chromatin dynamics

As chromatin actively interacts with the nuclear envelope^39^, the dynamic remodeling of the nucleus on viscoelastic substrates may influence chromatin dynamics. Given previous reports that cells lacking lamin A had increased chromatin dynamics in the nuclear interior^40,41^ and our observations that there was reduced lamin A/C protein expression and dynamic deformation of the nucleus on viscoelastic substrates (**Fig. 4**), we hypothesized that 2-kPa viscoelastic substrates promote chromatin dynamics in fibroblasts compared to 2-kPa elastic substrates. To test this, we stained fibroblasts on gel substrates with Hoechst dye to fluorescently label the DNA in cell nuclei and applied fluorescence recovery after bleaching (FRAP) techniques to observe the fluorescence recovery in euchromatic and heterochromatin regions. Immunofluorescence analysis of Hoechst-labeled fibroblasts with a heterochromatic histone mark (tri-methylated histone H3 on lysine 9, H3K9me3) or euchromatic mark (histone H3 acetylation AcH3), respectively, validated that in Hoechst-stained cell nuclei, heterochromatin appeared as the intensely stained regions that were perfectly colocalized with H3K9me3, and that euchromatin resided in the lightly stained regions as indicated by AcH3 signals (**Supplementary Fig. 11a-b**). These findings were further confirmed by co-staining fibroblasts expressing GFP-tagged HP1α (heterochromatin protein 1α) with Hoechst and AcH3 (**Supplementary Fig. 11c**), as HP1α is known as a major component of heterochromatin^42^.

Upon validating that Hoechst is a suitable candidate for FRAP experiments, FRAP results showed that the recovery half time (t_1/2_, the time taken for the bleached region to reach half of its final intensity) was shorter for cells on viscoelastic (slow-relaxing and fast-relaxing) substrates in both euchromatic and heterochromatic regions compared to elastic substrates, and no difference was seen between slow- and fast-relaxing substrates (**Fig. 5a-d**, **Supplementary Fig. 11d-e**, **Supplementary Videos 9-10**), which indicated that chromatin moves at a faster rate on viscoelastic substrates regardless of stress relaxation time. Moreover, to study how substrate viscoelasticity impacts the movement of nucleoproteins, we performed similar FRAP experiments utilizing fibroblasts expressing GFP-tagged HP1α (HP1α-GFP) cultured on substrates with different mechanical properties. Consistently, we found that the fluorescence in the bleached regions recovered faster on slow- and fast-relaxing substrates compared to elastic substrates (**Fig. 5e-f**, **Supplementary Fig. 11f**, **Supplementary Video 11**).

**Figure 5.**
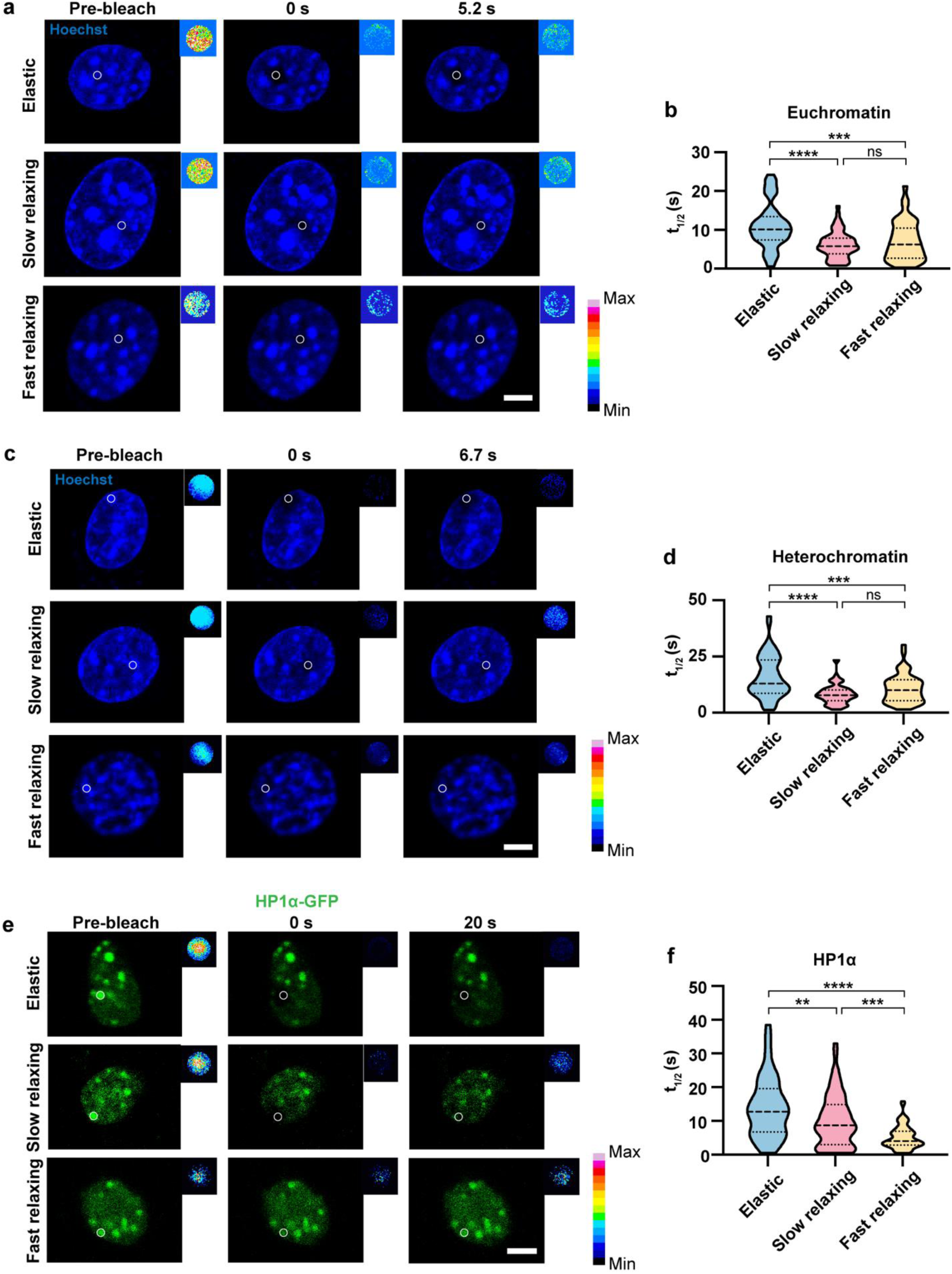
Viscoelastic substrates facilitate chromatin dynamics. **a-b**, Fluorescence recovery images of euchromatic regions after photo-bleaching in cells cultured on 2-kPa gels for 24 hours (**a**) and the truncated violin plots showing the recovery half time (t_1/2_, the time taken for the bleached region to reach half of its final intensity) of photo-bleached areas (white circle) (**b**) (n = 54, 56, 43 cells per condition from 3 independent experiments). Scale bar, 5 *μ*m. **c-d**, Fluorescence recovery images of heterochromatic regions after photo-bleaching in cells cultured on 2-kPa gels for 24 hours (**c**) and the truncated violin plots showing the recovery half time of photo-bleached areas (white circle) (**d**) (n = 57, 72, 51 cells per condition from 3 independent experiments). Scale bar, 5 *μ*m. **e-f**, Fluorescence recovery images after photo-bleaching in fibroblasts expressing GFP-tagged HP1α cultured on 2-kPa gels for 24 hours (**e**) and the truncated violin plots showing the recovery half time of photo-bleached areas (white circle) (**f**) (n = 56, 67, 68 cells per condition from 3 independent experiments). Scale bar, 5 *μ*m. Squares on the top right corner of each image indicates the fluorescence intensity in the photo-bleached area. In **b**, **d**, and **f**, the violins are drawn with the ends at the quartiles and the median as a horizontal line in the violin, where the curve of the violin extends to the minimum and maximum values in the data set. Significance was determined by one-way ANOVA using Tukey’s correction for multiple comparisons (\*\**P* < 0.01, \*\*\**P* < 0.001, \*\*\*\**P* < 0.0001).

The nucleolus is the largest subcompartment in the cell nucleus, which has important roles in ribosome biogenesis and chromatin architecture modulation^43^. To obtain a comprehensive landscape of chromatin dynamics, fibroblasts were transfected with a GFP-tagged fibrillarin plasmid and cultured on various substrates to track the movement of the nucleolus, as fibrillarin localizes to the dense fibrillar component of the nucleolus^44^. After recording the trajectories of fluorescently-labeled fibrillarin using confocal microscopy, we calculated the time-averaged mean square displacement (MSD) of the fibrillarin (**Supplementary Fig. 11g**). Interestingly, cells on purely elastic gels displayed a larger diffusion coefficient *D* of fibrillarin, but the approximate time for fibrillarin to diffuse 1 µm^2^ was similar between no- and slow-relaxing gels due to the larger diffusion exponent α on slow-relaxing gels (**Supplementary Fig. 11h-i**), suggesting that nucleolus movement is not sensitive to substrate viscoelasticity. Collectively, these results indicate that viscoelastic substrates enhance the motion of chromatin and some nucleoproteins in the cell nucleus, contributing to chromatin dynamics.

### Viscoelastic substrates enhance open chromatin marks

The decrease of chromatin condensation and increase in nuclear lamina remodeling and chromatin dynamics on soft viscoelastic substrates suggest an active remodeling of the chromatin, which may facilitate epigenetic changes and cell reprogramming in the presence of extrinsic factors (e.g., reprogramming transcriptional factors and cocktails of biochemical signals, etc.). To determine whether these physical changes are accompanied by biochemical epigenetic modifications of the chromatin, we examined the levels of histone marks associated with euchromatin and heterochromatin regions in the cell nucleus on 2-kPa elastic and slow-relaxing (viscoelastic) gels, as the chromatin dynamics was enhanced in cells cultured on slow-relaxing and fast-relaxing substrates to a similar extent.

Immunofluorescence analysis results revealed that several euchromatic histone marks, including AcH3, histone H3 lysine 27 acetylation (H3K27ac), and tri-methylated histone H3 on lysine 4 (H3K4me3), were significantly increased in cells on viscoelastic surfaces compared to elastic surfaces (**Fig. 6a-c, Supplementary Fig. 12a-c**), while we did not observe significant changes in heterochromatic marks between elastic and viscoelastic substrates, such as H3K9me3, and tri-methylated histone H3 on lysine 27 (H3K27me3) (**Supplementary Fig. 12d-e**). The increase in euchromatin marks was further confirmed by Western blotting (**Supplementary Fig. 12f-h**).

**Figure 6.**
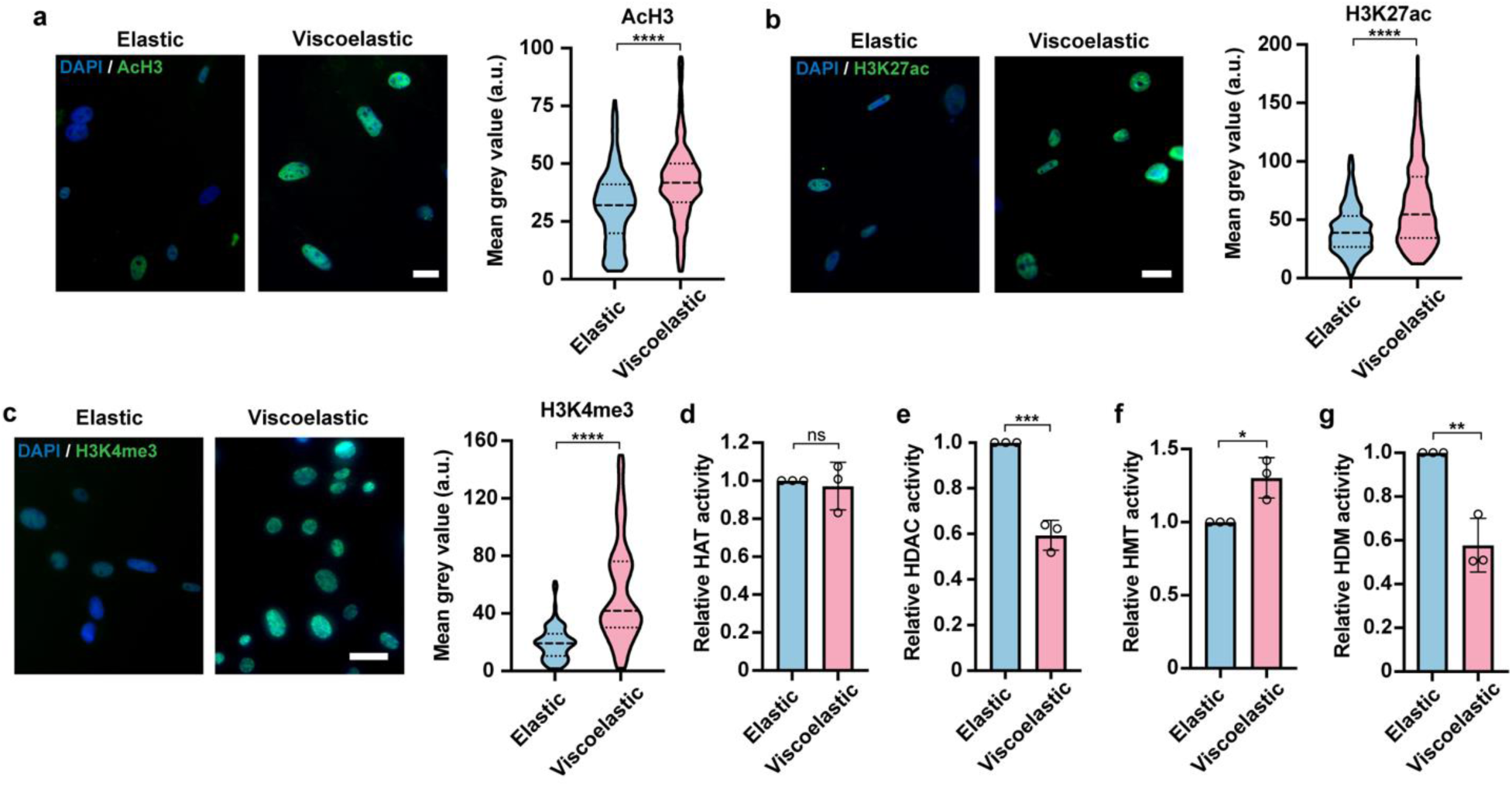
Viscoelastic substrates promote open chromatin structure. **a-c**, Fluorescent images (left) of histone H3 acetylation (AcH3, **a**), histone H3 lysine 27 acetylation (H3K27ac, **b**), and tri-methylated histone H3 on lysine 4 (H3K4me3, **c**) in mouse fibroblasts cultured on elastic or viscoelastic (slow relaxing) substrates of 2 kPa for 48 hours and truncated violin plots showing the fluorescence intensity quantification (right) (n = 342, 302 cells from 3 independent experiments per condition for AcH3; 468, 626 cells from 3 independent experiments per condition for H3K27ac; 250, 180 cells from 3 independent experiments per condition for H3K4me3). Scale bar, 20 *μ*m. **d-e**, Quantification of histone acetyltransferase (HAT) activity (**d**) and histone deacetylase (HDAC) activity (**e**) in cells cultured on elastic or viscoelastic gels for 48 hours (n = 3 independent experiments, 500,000 cells were initially seeded). **f-g**, Quantification of H3K4-specific histone methyltransferase (HMT) activity (**f**) and histone demethylase (HDM) activity (**g**) in cells cultured on elastic or viscoelastic substrates for 48 hours (n = 3 independent experiments, 500,000 cells were initially seeded). For truncated violin plots in **a-c**, the violins are drawn with the ends at the quartiles and the median as a horizontal line in the violin, where the curve of the violin extends to the minimum and maximum values in the data set. In **d-g**, bar graphs show the mean ± s.d. A two-tailed, unpaired t test was used to determine the statistical significance (\**P* < 0.05, \*\**P* < 0.01, \*\*\**P* < 0.001, \*\*\*\**P* < 0.0001).

Next, we analyzed the activities of chromatin-modifying enzymes to investigate their contributions to these epigenetic alterations. Though there was no significant change in histone acetyltransferase (HAT) activity in fibroblasts on viscoelastic gels, we found that histone deacetylase (HDAC) activity in cells on viscoelastic surfaces was reduced by half (**Fig. 6d-e**), which could lead to the elevated acetylation level on histone H3. In addition, viscoelastic substrates increased the activity of H3K4-specific histone methyltransferase (HMT) and decreased demethylase (HDM) activity (**Fig. 6f-g**), which could enhance H3K4me3 in cells on viscoelastic surfaces. These findings suggest that viscoelastic substrates regulate histone-modifying enzymes (HDAC, H3K4-specific HMT, and H3K4-specific HDM) that work together to modulate the epigenetic state of cells.

### Viscoelastic substrates promote chromatin accessibility in Ascl1- and Oct4-targeting genomic regions

Our findings demonstrate that substrate viscoelasticity promotes physical remodeling of nuclear structures and biochemical modification-associated nuclear events towards a more open chromatin architecture and enhanced chromatin dynamics (**Fig. 1h-i**, **Supplementary Fig. 3c**, **Fig. 5, Supplementary Fig. 11**). Moreover, cells on viscoelastic substrates had increased expression of euchromatic marks (**Fig. 6a-c, Supplementary Fig. 12a-c**), potentially leading to a more open chromatin structure. Consequently, we hypothesized that fibroblasts on elastic and viscoelastic substrates may have distinct chromatin accessibility landscapes, where different combinations of permissible chromatin regions can lead to active or silenced transcriptomic activities in response to external perturbations. To investigate this, we performed assay for transposase-accessible chromatin with sequencing (ATAC-seq) on fibroblasts cultured on elastic and slow-relaxing (viscoelastic) substrates of 2 kPa to investigate the influence of substrate viscoelasticity on genome-wide chromatin accessibility at the population level.

PCA results of ATAC-seq signals placed elastic groups away from viscoelastic groups along PC1 that captured 97% of variance, which demonstrated that substrate viscoelasticity altered accessible chromatin regions across the genome (**Fig. 7a**). By differential peak analysis, surprisingly, we found ∼40 % of peaks were up-regulated to have higher chromatin accessibility in viscoelastic groups compared to elastic groups (**Supplementary Fig. 13a-c**). We then annotated the differentially expressed peaks to associate them with genomic features including nearest genes and *cis*-regulatory elements such as promoters and enhancers. Compared to the genomic distribution of down-regulated peaks, a higher percentage of the up-regulated peaks on viscoelastic surfaces (11.7 %) were located in promoter regions in comparison to elastic surfaces (6.9 %) (**Fig. 7b**). Since promoters are the initiation point of transcription, this shift suggested that viscoelastic substrates may lower the epigenetic barrier for induced gene expression by external transcription factors.

**Figure 7.**
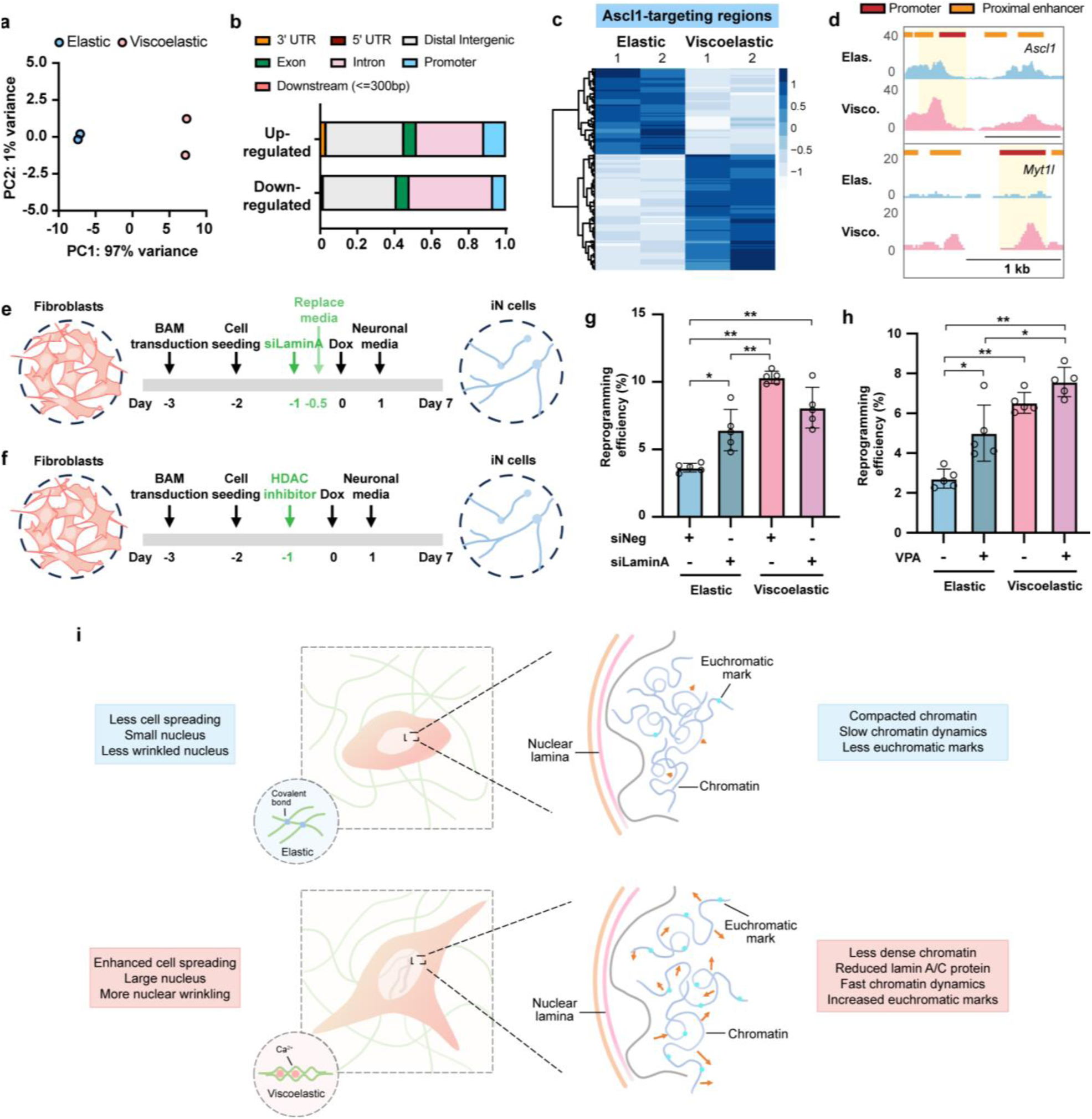
Viscoelastic substrates regulate genome-wide chromatin accessibility. **a**, Principal component analysis (PCA) of ATAC-seq data from independent biological replicates of mouse fibroblasts cultured on 2-kPa elastic or viscoelastic (slow relaxing) substrates for 48 hours (n = 2 biological replicates). **b**, Percentage of genomic features associated with up- or down-regulated ATAC peaks in viscoelastic groups in comparison with elastic groups. **c**, Heatmaps showing the differentially expressed ATAC signals in Ascl1-targeting genomic sites. **d**, ATAC peaks around the promoters of *Ascl1* and *Myt1l* in fibroblasts cultured on elastic and viscoelastic gels. **e-f**, Schematic showing the timeline of *Lmna* knock-down (**e**) or HDAC inhibition (**f**) for reprogramming experiments. **g**, Quantification of the reprogramming efficiency of fibroblasts transfected with negative control or *Lmna* siRNA and cultured on various surfaces for 7 days (n = 5 independent experiments). **h**, Quantification of the reprogramming efficiency of fibroblasts cultured on various surfaces and in the absence or presence of an HDAC inhibitor (valproic acid, VPA, 2.5 mg ml^−1^) (n = 5 independent experiments). **i**, Schematic illustrating the effects of viscoelasticity on the cell nucleus. In **g** and **h**, bar graphs show the mean ± s.d. and a two-way ANOVA using Tukey’s correction for multiple comparisons was used to determine the significance between the groups (\**P* < 0.05, \*\**P* < 0.01).

To further test whether substrate viscoelasticity can assist lineage-specific cell phenotype conversion and reprogramming, we examined chromatin accessibility around lineage-determining transcription factors in elastic and viscoelastic groups using publicly available chromatin immunoprecipitation followed by sequencing (ChIP-seq) datasets. By plotting differentially expressed ATAC signals in genomic regions interacting with certain transcription factors identified from their ChIP-seq datasets, we were able to explore the accessibility changes around the genomic targets of pioneer transcription factors that play important roles in regulating cell fate decisions in development and cell reprogramming. Analysis of Ascl1 (neuronal lineage)^33^, MyoD (myogenic lineage)^45^, ETV2 (endothelial and hematopoietic lineage)^46^, and Oct4 (pluripotency)^47^ target sites demonstrated that substrate viscoelasticity remodeled the chromatin to increase its accessibility around genomic regions actively involved in neuronal and pluripotent lineages (**Fig. 7c**, **Supplementary Fig. 13d**), while regions relevant to the hematoendothelial lineage tended to be closed on viscoelastic surfaces (**Supplementary Fig. 13e-f**). For example, the promoter regions of *Ascl1* and *Myt1l* were more permissible on viscoelastic substrates (**Fig. 7d**), making it easier for fibroblasts cultured on viscoelastic substrates to activate the endogenous expression of these two genes in cell reprogramming and facilitate the cell phenotype conversion. This was supported by the increased gene expression of endogenous Ascl1 on slow-relaxing substrates compared to elastic gels (**Fig. 3d**). In addition, the promoters of typical mesenchymal cell markers (*Eng* [CD105] and *Nt5e* [CD73]) and an epithelial-mesenchymal transition regulator (*Prrx1*) displayed lower accessibility in viscoelastic groups compared to elastic groups (**Supplementary Fig. 13g**), implicating that viscoelastic substrates lower the mesenchymal barrier in fibroblasts to undergo cell reprogramming.

### Substrate viscoelasticity regulates cell reprogramming through epigenetic remodeling

To directly determine the roles of viscoelastic substrate-induced epigenetic changes such as the downregulation of lamin A and the decrease of HDAC activity in cell reprogramming, we knocked down *Lmna* and inhibited HDAC, respectively, and analyzed iN reprogramming efficiency (**Fig. 7e-f**).

By transfecting BAM-transduced fibroblasts on elastic and viscoelastic substrates with a small interfering RNA (siRNA), we silenced *Lmna* to determine whether the nuclear lamina was involved in viscoelasticity-enhanced iN reprogramming (**Fig. 7e, Supplementary Fig. 14a-b**). We found that there was an increase in the reprogramming efficiency in cells on elastic surfaces where *Lmna* was silenced, compared to non-silenced cells, but the iN reprogramming facilitated by siRNA did not further improve the efficiency on a viscoelastic substrate (**Fig. 7g**). Additionally, a higher proportion of fibroblasts were reprogrammed into iN cells on glass when *Lmna* was knocked down (**Supplementary Fig. 14c**), though this was still not as efficient as cells on viscoelastic substrates. It is worth noting that the reduction of A-type lamins by siRNA was much higher than the intrinsic lamin A/C decrease induced by cell nuclear sensing of substrate viscoelasticity (**Fig. 4a-b**, **Supplementary Fig. 14a-b**). Since loss of lamin A impairs nuclear stability and integrity in many cases^48–50^, exaggerated silencing of *Lmna* may not be the optimal choice to boost cell reprogramming. Together, these results suggest that lamin A mediates the effects of substrate viscoelasticity on the cell reprogramming process.

Next, valproic acid (VPA) was employed to inhibit HDAC function, resulting in an increase in histone acetylation and thus, contributes to a more open chromatin structure (**Fig. 7f**, **Supplementary Fig. 14d-e**). HDAC inhibition enhanced the iN reprogramming efficiency of cells on elastic substrates to an extent similar to cells on viscoelastic substrates without inhibitor treatment, but did not further enhance the reprogramming efficiency on viscoelastic surfaces (**Fig. 7h**). The reprogramming efficiency on glass was also elevated with the treatment of VPA, but to a much lesser extent than viscoelastic substrates (**Supplementary Fig. 14f**). These findings reflect that the downregulation of HDAC by viscoelastic substrates may play a role in regulating viscoelasticity-mediated cell phenotype conversion.

## Outlook

Here, we demonstrate, for the first time, that cell nucleus is sensitive to substrate viscoelasticity, and viscoelastic substrates induce a more ‘plastic’ epigenetic state of adherent cells that facilitates cell reprogramming. This includes reduced lamin A/C expression, increased nuclear lamina wrinkling and deformation, decreased chromatin condensation, faster chromatin dynamics, enhanced euchromatin marks, and greater chromatin accessibility at specific genomic regions, all of which are more pronounced on softer (2 kPa) viscoelastic substrates (**Fig. 7i**). On stiffer surfaces (e.g., 20 kPa), enhanced cell adhesion results in fully spread cells, which dominates the mechanotransduction response and may overshadow the effects of viscoelasticity. For instance, only fast-relaxing gels of 20 kPa are associated with an increase in nuclear volume. It is also worth noting that substrate viscoelasticity modulates gene expression profiles in a manner distinct from substrate stiffness; furthermore, substrate viscoelasticity reduces the sensitivity of gene expression profiles to variations in stiffness (**Fig. 2**). These findings demonstrate that cells can distinguish between the effects of substrate viscoelasticity and stiffness, likely through temporal responses, such as relaxation, creep, and the dynamic behavior of both the substrate and intracellular force-generating structures, including focal adhesions, the cytoskeleton, and the nuclear lamina. Conversely, the overlapping intracellular mechanosensing and mechanotransduction components for substrate viscoelasticity and stiffness may explain the ‘masking’ effect of viscoelasticity on stiffness, warranting further investigation.

Our studies demonstrate three potential mechanisms of substrate viscoelasticity-mediated nuclear remodeling, where more pronounced effects were observed on soft surfaces. First, there is a direct mechanical coupling by which viscoelastic substrates modulate the physical properties of the cell nucleus. More compliant viscoelastic substrates (2 kPa) induce a less compacted chromatin architecture and faster chromatin dynamics, which may be linked to dynamic changes in focal adhesions, cytoskeletal reorganization, and nuclear lamina structure. Although soft substrates can only sustain low levels of traction forces exerted by cells^51,52^, the stress relaxation and creep due to viscoelasticity may facilitate more dynamic force generation by the actin cytoskeleton and allow sufficient time for focal adhesion assembly, which results in increased cell spreading^6^ and a larger nuclear volume (**Fig. 1d-g**). The dynamic redistribution of intracellular forces caused by viscoelastic surfaces, along with reduced cytoskeletal constraints around the nucleus on soft surfaces^40,53,54^, may lead to increased nuclear wrinkling and deformation (**Fig. 4e-g**), which in turn results in less compacted and more dynamic chromatin (**Fig. 1h-i**, **Fig. 5**).

Secondly, the reduction of lamin A/C protein expression in cells grown on viscoelastic substrates represents another critical mechanism of mechanosensing. It is well-established that lamin expression scales with matrix stiffness^55^. Our RNA-seq data indicates that the viscoelasticity of substrates (both 2 and 20 kPa) has a more pronounced effect on lamin A/C downregulation than purely soft elastic surfaces (2 kPa compared to a 20 kPa elastic surface) (**Fig. 2e**). Furthermore, lamin knockdown on soft surfaces enhances iN reprogramming efficiency, partially mimicking the effects of substrate viscoelasticity (**Fig. 7e, g**), suggesting that lamin A/C plays a key role in the nuclear sensing of substrate viscoelasticity to facilitate chromatin and epigenome remodeling for cell reprogramming. It has been reported that lamin A/C-deficient fibroblasts are not able to form the perinuclear apical actin cables that protect the cell nucleus from nuclear deformation^56^, which could contribute to increased nuclear deformation, enhanced nuclear movement, less compact chromatin, and faster chromatin dynamics. Notably, the chromatin landscapes modulated by viscoelastic substrates resemble those of undifferentiated embryonic stem cells (ESCs) that lack lamin A/C and exhibit minimal nuclear lamina constraints, decondensed chromatin organization, enhanced chromatin protein dynamics, and a higher enrichment in euchromatic marks compared to differentiated cells^37,57–59^. This unique chromatin state renders ESCs highly plastic and capable of differentiating into various cell types. Our findings suggest that substrate viscoelasticity may promote cellular plasticity in a manner similar to that of stem cells, thereby creating a favorable epigenetic landscape for cell phenotype transitions. However, the mechanisms by which substrate viscoelasticity and stiffness regulate lamin expression remain to be elucidated in future studies.

Thirdly, viscoelastic substrates can regulate epigenetic marks through mechano-chemical signaling involving epigenome-modifying enzymes. Viscoelastic substrates decrease HDAC activity, enhance H3K4-specific HMT activity, and decrease H3K4-specific HDM activity, collectively resulting in elevated levels of euchromatin marks such as AcH3, H3K27ac, and H3K4me3 (**Fig. 6**). These changes may contribute to the improved efficiency of cell reprogramming and are consistent with increased chromatin accessibility at the promoters and enhancers of neuronal and pluripotent genes. In line with our findings, previous studies have demonstrated that HDAC inhibition can enhance cell reprogramming efficiency^60–62^. Given that several HDAC subtypes are known to participate in genome-nuclear lamina interactions to regulate cell fate decisions^63–65^, there may be an interplay between nuclear lamina dynamics and HDAC activity that contributes to enhanced cellular plasticity on viscoelastic substrates. This highlights the need for further investigation into the roles of specific HDAC subtypes in response to substrate viscoelasticity-mediated nuclear lamina remodeling. Additionally, other biochemical signaling pathways for epigenetic regulation such as histone methylation and other modifications may also contribute to the viscoelasticity-enhanced cell reprogramming process. For instance, our RNA-seq data revealed the potential involvement of other chromatin modifications, such as ADP-ribosylation and sumoylation, as *Parp1* and *Sumo1* genes were differentially expressed on elastic versus viscoelastic substrates (**Fig. 2f, Supplementary Fig. 6b**). It is also possible that non-coding RNAs play a role in the modulation of cell phenotype changes through their buffering abilities towards the expression level of nucleoskeletal, cytoskeletal, and ECM proteins^66–68^. Additionally, as it has been reported that matrix viscoelasticity promotes neural maturation of 3D-encapsulated human iPSC-induced neural cells^69^, it would be interesting to explore the viscoelastic effects on cell reprogramming in 3D contexts in future studies.

As marked alterations in tissue mechanics occur enormously in development, aging, and disease progression^70–72^, our findings of the stiffness-dependent viscoelastic effects on the cell nucleus, chromatin, and the epigenome will have important implications in cellular responses to the changing ECM mechanical properties during aging and disease development. For example, diverse pathological conditions are accompanied with an increase in viscoelasticity and stiffness, such as fibrosis^73^ and most malignant tumors^74^, while some diseases (e.g., diabetes and obesity) can make the local tissue softer and more viscoelastic. These changes in mechanical environment may have profound effects on the differentiation, de-differentiation and trans-differentiation of tissue-resident cells, which warrants further investigations in disease-specific context.

## Methods

All experiments were performed in accordance with relevant guidelines and ethical regulations approved by the UCLA Institutional Biosafety Committee (BUA-2016-222).

### Alginate hydrogel preparation

Sodium alginate (Sigma-Aldrich) with over 60% of its unit as guluronate monomer units and an average molecular weight over 200 kDa (PRONOVA UP MVG, viscosity > 200 mPa s) was used to prepare elastic and slow-relaxing (viscoelastic) hydrogels. Sodium alginate with a very low viscosity (PRONOVA UP VLVG, M_w_ < 75 kDa, viscosity < 20 mPa s) was mixed with MVG alginate at a ratio of 2:1 to fabricate the fast-relaxing (viscoelastic) hydrogels. Briefly, sodium alginate was dissolved in MES buffer (0.1 M MES, 0.3 M NaCl, pH 6.5) or Dulbecco’s modified Eagle’s medium (DMEM) at 2.5% wt/vol to make the alginate solution for elastic or viscoelastic gels. Crosslinker solutions was prepared prior to gel fabrication, where appropriate amounts of adipic acid dihydrazide (AAD, from Sigma) was dissolved in MES buffer containing 2.5 mg ml^−1^ of 1-Hydroxybenzotriazole (HOBt, from Sigma) and 50 mg ml^−1^ N-(3-Dimethylaminopropyl)-N’-ethylcarbodiimide (EDC, from Sigma) as covalent crosslinker solutions and different concentrations of calcium sulfate was evenly distributed into DMEM as ionic crosslinker solutions. 0.8 ml of alginate solution was mixed with 0.2 ml of crosslinker solution using two syringes and deposited between two hydrophobically treated glass slides with a spacer of various heights. Ionically crosslinked hydrogels and covalently crosslinked hydrogels were put into DMEM after 2 or 12 hours of gelation, respectively, and washed two times per day for 2 days to remove unused crosslinkers. For cell-adhesive hydrogels, an oligopeptide with the sequence of GGGGRGDSP (Genscript) of 100 mg was conjugated to the alginate polymer (1 g) through the reaction with EDC (484 mg) and N-hydroxysulfosuccinimide (sulfo-NHS, 274 mg) in 1% (wt/vol in MES buffer) alginate solution for 20 h. The resulting solution was dialyzed against deionized water for 3 days (molecular weight cutoff of 3.5 kDa), sterile-filtered, and lyophilized before being reconstituted into MES buffer or DMEM to make the hydrogels. The concentrations of AAD solutions to make elastic gels were 2, 6, and 15 mM, while the calcium concentrations in the crosslinker solutions used to fabricate viscoelastic gels were 35, 90, and 180 mM for slow-relaxing gels and 55, 120, and 300 mM for fast-relaxing gels, respectively. The final concentration of alginate in the hydrogels was 2% (wt/vol). On average, there were ∼62 RGD peptides per MVG alginate chain (**Supplementary Note 1**, **Supplementary Fig. 15-16**). Since the same mass amount of alginate and peptide was used in each conjugation reaction, the ligand density among various gels should be identical.

### Quantitative nuclear magnetic resonance

To determine the average ligand density of polymer chains in alginate-based hydrogels, NMR solutions (20 mg of unconjugated or RGD-coupled sodium alginate per 1 ml of D_2_O) were prepared and heated overnight at 40^°^C to ensure complete dissolution of the alginate. A solution of maleic acid NMR standard (4 mg of standard per 1 ml of D_2_O) was prepared and transferred to a sealed capillary tube to prevent precipitation of the alginate. NMR spectra were recorded on a Bruker DRX500 spectrometer at 383 K in D_2_O. Chemical shifts were reported with respect to the solvent residual peak at 4.79 ppm.

### Characterization of gel swelling

Alginate hydrogels were fabricated and incubated in DMEM at 37 °C for 1 or 7 days. The gels were then taken from the medium and their diameter and height was measured using a caliper to calculate the volume. Gel swelling was quantified as the ratio of the volume of gels at day 1 or day 7 to their volumes before immersion.

### Mechanical characterization of hydrogels

The mechanical properties of covalently and ionically crosslinked hydrogels were determined using rheology measurements and compression tests. To measure the rheological properties, hydrogels were made on the same day of the rheology test. A small-amplitude strain sweep with a constant frequency at 0.5 rad/s was performed to find the linear elastic regime, followed by a frequency sweep at 0.5% strain and a full-amplitude strain sweep at 0.5 rad/s to assess the storage modulus and loss modulus using a rheometer (Anton Paar MCR302). The stiffness and stress relaxation properties were measured from compression tests on alginate gels after one-day equilibrium in DMEM. The gels were compressed to 15% strain with a deformation rate of 1 mm min^−1^ using a Chatillon TCD 225 series Force Measurement System. Afterward, the strain was held constant and the load was recorded over time. The initial stiffness of the hydrogels was calculated using the slope of the stress-strain curve at 5-10% strain. To find *τ*_½_ (the time that gels took to relax their initial stress to its half value) of different gels, stress-relaxing curves were fitted into a biexponential equation based on a two-element Maxwell-Weichert model and *τ*_½_ was calculated from each curve.

### Characterization of gel degradation

Alginate hydrogels were fabricated and incubated in DMEM at 37 °C for 1 or 7 days. The gels were then removed from the medium, frozen at −80 °C, and lyophilized. The dry mass of the hydrogels at day 1, day 7 and before incubation was measured following lyophilization. The weight ratio of the dry mass of gels after 1 or 7 days of DMEM immersion to the dry mass of gels before immersion was calculated.

### Fibroblast isolation, culture, and cell seeding

Mice utilized in these studies were housed under specific pathogen-free conditions and 12-hour-light/12-hour-dark cycles with a control of temperature (20-26 °C) and humidity (30-70%). All experiments, including breeding, maintenance and euthanasia of animals, were performed in accordance with relevant guidelines and ethical regulations approved by the UCLA Institutional Animal Care and Use Committee (protocol no. ARC-2016-036 and ARC-2016-101).

Ear fibroblasts were isolated from adult C57BL/6 mice of 1 month old (male and female, from Jackson Laboratory, 000664) and Tau-EGFP reporter mice (Jackson Laboratory, 004779). Cells were expanded in mouse embryonic fibroblast (MEF) medium after isolation: DMEM (Gibco) with the addition of 10% fetal bovine serum (FBS, from Gibco) and 1% penicillin/streptomycin (Gibco), and used at passage 2 for all experiments. Fibroblasts were plated on the surface of hydrogels at an initial density of 10,000 cells cm^−2^ to allow for cell-ECM interactions with minimal interference from cell-cell interactions. MEF medium was added to the hydrogels 2 hours after cell seeding to ensure cell attachment. To observe cell spreading and cell nuclear differences on different substrates, cells were cultured on hydrogels for 24 hours before being fixed and stained for analysis.

### Cell viability assay

Live and dead assays were performed on cells cultured on various substrates and tissue culture plates for 24, 48, and 72 hours using the LIVE/DEAD Cell Imaging Kit (Invitrogen, R37601) according to the manufacturer’s protocol. Cells were incubated with an equal volume of ×2 working solution for 15 minutes at room temperature. Epifluorescence images were collected using a Zeiss Axio Observer Z1 inverted fluorescence microscope and analysed using Fiji.

### EdU labeling and Staining

EdU labeling and staining was performed on cells culture on various substrates for 48 hours using the Click-iT EdU Alexa Fluor 488 Imaging Kit (ThermoFisher, C10337) according to the manufacturer’s protocol. Briefly, fibroblasts were incubated with 10 μM EdU for 2 hours before sample collection. Samples were fixed with 4% paraformaldehyde (Electron Microscopy Sciences, 15710) for 15 minutes at room temperature, washed twice with 3% bovine serum albumin (BSA; Fisher, BP1600) in serum-free DMEM and permeabilized with 0.5% Triton X-100 (Sigma, T8787) in serum-free DMEM for 20 minutes. Then, samples were washed twice with 3% BSA followed by incubation with the EdU cocktail reaction for 30 minutes. After washing twice with 3% BSA, samples were stained with 4,6-diamino-2-phenylindole (DAPI; Invitrogen, D3571) for 10 minutes to identify nuclei.

### Viral production and cell transduction

Dox-inducible lentiviral vectors for Tet-O-FUW-Ascl1, Tet-O-FUW-Brn2, Tet-O-FUW-Myt1l, Tet-O-FUW-GFP, and FUW-rtTA plasmids were used to transduce fibroblasts for ectopic expression of Ascl1, Brn2, Myt1l, GFP, and reverse tetracycline transactivator (rtTA). The STEMCCA lentiviral vector was used for the ectopic expression of OSKM^12^. The pBABE-puro-GFP-wt-Lamin A retroviral vector (AddGene, 17662) was used for the ectopic expression of lamin A-GFP^12^. Lentivirus was produced using established calcium phosphate transfection methods, while retrovirus was produced using Platinum E cells and FuGENE HD (Promega, E2311) as previously described^75^. Viral particles were collected and concentrated using Lenti-X Concentrator (Clontech) according to the manufacturer’s protocol. Stable virus was aliquoted and stored at - 80°C. For viral transduction, fibroblasts were seeded and allowed to attach overnight before incubation with the virus and polybrene (8 µg ml^−1^, Sigma) for 24 h. After incubation, transduced cells were seeded onto hydrogels.

### Direct reprogramming of fibroblasts into induced neuronal cells

Fibroblasts were transduced with lentivirus encoding BAM a day before being seeded onto hydrogels. Cells were then seeded on various substrates with the same initial cell density of 10,000 cells cm^−2^. After 24 h of cell seeding, the medium was changed to MEF medium containing doxycycline (Dox, 2 µg ml^−1^, from Sigma) to activate transgene expression (day 0). On day 1, the medium was replaced with N3 medium: DMEM/F12 (Gibco) with the addition of N2 supplement (Invitrogen), B27 supplement (Invitrogen), 1% penicillin/streptomycin (Gibco), and Dox (2 µg ml^−1^, from Sigma). Culture medium was replenished every other day during reprogramming to maintain the activity of the supplements. Cells were collected at day 2, 5, 7, 21, and 28 and analyzed to monitor the viscoelastic effects on cell phenotype conversion during the course of iN reprogramming. To quantify the reprogramming efficiency from fibroblasts to induced neuronal (iN) cells, the samples were fixed and immunostained for Tubb3 (neuron-specific class III beta-tubulin) after 7 days of culturing. iNs were identified based on displays of a typical neuronal morphology (defined as cells with a circular cell body containing a neurite that was at least two times the length of the cell body) and positive Tubb3 expression as previously described^32^. The reprogramming efficiency was calculated as the percentage of Tubb3^+^ iN cells from the total cells initially plated.

### Direct reprogramming of fibroblasts into induced pluripotent stem cells (iPSCs) and *in vitro* differentiation of iPSCs

Fibroblasts were transduced with lentivirus encoding OSKM a day before being seeded onto hydrogels. With the initial cell number of 5,000 per well in a 24-well plate, cells were kept in MEF medium for 24 h and then the medium was changed to mouse embryonic stem cell (mESC) medium: DMEM with the addition of 15% mouse ESC maintenance FBS (STEMCELL Technologies), 100 units ml^−1^ ESGRO mouse leukaemia inhibitory factor (mLIF, from Millipore), 0.1 mM β-mercaptoethanol (Sigma) and 1× MEM non-essential amino acid (Gibco). The medium was replenished every day for 10 days. To count iPSC colonies, samples were fixed and immunostained for Nanog after 10 days of culturing. Colonies with positive Nanog expression were identified as iPSC colonies.

After 10 days of reprogramming, iPSC colonies were selected to form embryoid bodies for *in vitro* differentiation using the hanging-drop method in mESC maintenance media without mLIF. Embryoid bodies were then plated onto gelatin-coated surfaces and cultured under distinct differentiation media to promote differentiation into various cell types. After 2 weeks of differentiation, samples were fixed and immunostained to determine the derived cell phenotypes.

### Immunofluorescence staining

For immunostaining, cells on various substrates were fixed with 4% paraformaldehyde (Electron Microscopy Sciences) in serum-free DMEM for 15 min after removing the media and washing with serum-free DMEM once. Samples were then washed three times in serum-free DMEM, permeabilized with 0.5% Triton-X-100 (Sigma), and blocked with 5% donkey serum (Jackson Immunoresearch) in serum-free DMEM following standard immunostaining protocols^34^. For actin-cytoskeleton staining, samples were incubated with Alexa Fluor 635 phalloidin (Invitrogen) for 1 h. Primary antibodies (refer to **Table S1, Supplementary Information**) were incubated overnight at 4 °C, followed by 1-h incubation with Alexa 488 and/or Alexa 546-labeled secondary antibodies (Invitrogen). Nuclei were stained with DAPI (Invitrogen). Epifluorescence images were collected using a Zeiss Axio Observer Z1 inverted fluorescence microscope, whereas confocal images were acquired using Leica SP8 Confocal Laser Scanning microscope. Fluorescence quantifications were performed using Fiji macros and/or CellProfiler pipelines and the methods are described in the corresponding sections.

### Quantification of cell spreading

To identify the boundary of fibroblasts, Cellpose^76^ was used to segment cell bodies from microscopy images showing DAPI and phalloidin staining. The mask files were then imported into CellProfiler to measure the area of each cell, where cells at image borders were excluded to achieve maximum accuracy.

### Quantification of nuclear volume and chromatin compaction

To quantify nuclear volume and chromatin compaction index, DAPI-stained nuclei images were taken from the bottom to the top using the z-stack function with a step size of 300 nm by a Leica SP8 Confocal Laser Scanning microscope. For each individual nuclei, the maximum intensity was projected using Fiji, followed by Gaussian blur and thresholding to obtain the mask. The mask was then applied to each individual slice of the corresponding nuclei to quantify the area of the nuclei sections. Nuclear volume was calculated as the sum of the area in each slice multiplied by the step size. Chromatin compaction index was determined as the ratio of the integrated fluorescence intensity of the sum intensity of all slices for each individual nuclei to the corresponding nuclear volume as previously described^29^.

### Quantification of lamin wrinkles

To quantify the degree of nuclear wrinkling indicated by nuclear lamin staining, lamin A/C and lamin B1 wrinkling indices were calculated based on the output from the FeatureJ Edge tracking algorithm (https://imagescience.org/meijering/software/featurej/) following a previously reported quantification method^77^. First, the edges of nuclei were determined from the maximum projection of the z-stacked lamin staining images by Gaussian blur and thresholding in Fiji. Secondly, the nucleoplasm boundaries were identified using the erode function on the identified cell nucleus borders. Same blurring parameter, thresholding selection, and erode parameter were kept identical across groups to make fair comparisons. Lastly, FeatureJ Edge contour tracking were applied in the maximum projection images to detect edges (wrinkles) and the mean intensity inside the nucleoplasm in the edge detection output images were measured as lamin wrinkling indices. Analysis confirmed that lamin wrinkling index did not correlate with nuclear area (i.e. large nuclei did not necessarily have more wrinkles than small nuclei)^77^ and fluorescence variation inside nucleoplasm (i.e. a no-wrinkle nucleus with high fluorescence intensity in its nucleoplasm will not obtain a large wrinkling index number, **Supplementary Fig. 17**).

### Fluorescence quantification of epigenetic marks

To measure fluorescence intensity of histone staining, DAPI-stained nuclei were segmented using Gaussian blur, thresholding, watershed, and analyze particle functions in Fiji to identify individual nuclei. The obtained masks were then applied to the corresponding stained fluorescence channel to quantify the average fluorescence intensity within each nucleus.

### RNA sequencing

500,000 fibroblasts were seeded on substrates with various stiffness and viscoelasticity for 48 hours and collected to isolate RNA and prepare the library, with the assistance of the Technology Center for Genomics & Bioinformatics at UCLA. Raw files were trimmed with Trimmomatic^78^ using default settings and then aligned to mouse reference genome (mm10) using STAR^79^ with default parameters. Transcriptome alignments were quantified using featureCounts^80^ with the GENCODE annotation file. Differentially expressed genes (DEGs) were identified with DESeq2^81^ using *p*_adj_ = 0.05 as the threshold. Gene ontology (GO) enrichment analysis was performed using the enrichGO function in the clusterProfiler package^82^. To build the mechanotransduction-related gene regulatory network database, Harmonizome^83^ was used to search functional genes by categories. For heatmap of DEGs, values were normalized using size factors estimated from gene count matrices by DESeq2 and then z-scored by row.

### Nuclear stiffness measurement

A nano-indentation platform (Pavone Nanoindenter, Optics11 Life) was employed to quantify the apparent Young’s modulus of the cell nucleus of fibroblasts on various gel substrates. A probe tip with a stiffness of 0.58 N/m and radius of 3.5 μm was calibrated in MEF culture medium with a geometrical factor of < 5 % variation than in air. Cell samples were placed in MEF medium for measurements. The Displacement Control mode indentation was first used to identify a proper probe force and the Peak Poking Control mode (max load 0.02 μN, piezo speed 10 μm s^−1^) was used for measurements. Nano-indentations were performed across pre-selected cells identified by image acquisition with probing locations/coordinates. The Hertzian model fitting to indentation profiling (up to 16% probe tip radius according to the Pavone instrument manual) was used to extract the Young’s modulus of samples.

### Fluorescence recovery after photobleaching (FRAP)

Fibroblasts were seeded on hydrogels for 24 h and stained with Hoechst 33342 (Invitrogen) for FRAP experiments using a Leica TCS SP8 microscope with a pinhole of 2.5 to capture more spatial information. Briefly, a prescan of 7 images was acquired with an interval of 740 ms, followed by a bleaching pulse of 20 iterations at the selected area of 1 μm in diameter. 60-180 images were then taken with an interval of 1 s to record the changing fluorescence over time. To engineer fibroblasts with the expression of GFP-tagged HP1α, fibroblasts were transfected with a HP1α-GFP plasmid (AddGene, 17652) using lipofectamine 3000. The media was changed after 12 hours of plasmid transfection. Twenty-four hours later, HP1α-GFP cells were seeded on various gels for 24 hours, followed by FRAP experiments. For HP1α-GFP fibroblasts, a prescan of 7 images was acquired with an interval of 740 ms, followed by a bleaching pulse of 10 iterations at the selected area of 1 μm in diameter. 60 images were then taken with an interval of 1 s to record the changing fluorescence over time. Analysis of FRAP movies were performed using a Jython script in Fiji.

### Fibrillarin mean square displacement (MSD)

Fibroblasts were transfected with a pCMV3-FBL-GFPSpark plasmid (MG53664-ACG, Sino Biological) using lipofectamine 3000. The media was changed to MEF media at 12 hours after plasmid transfection and the cells were allowed to grow for 12 hours before being plated on various substrates. Twenty-four hours after cell seeding, fibroblasts were stained for Hoechst 33342 (Invitrogen) for 10 minutes to label cell nuclei and imaged using a Leica TCS SP8 microscope in a live cell chamber (37°C, 5% CO_2_) for 20 time points with 2-min intervals. Z-stack images of Hoechst and GFP channels were taken to capture the potential stage drift and nucleus drift and rotation.

TrackMate in Fiji was employed to extract the movement trajectories of fibrillarin. Only trajectories with no concatenation between different loci were used in the analysis. Upon obtaining the trajectories, nucleus drift and rotation correction were performed to minimize artifacts following a previous report^40^. Briefly, the trajectory of the mass center of each cell nuclei was obtained from the Hoechst channel and subtracted from the fibrillarin trajectories to correct for nuclear drift. Then, the transformation matrix for each time point was extracted to calculate the average rotation matrix of each trajectory, and the trajectories were multiplied by their inverse rotation matrix to correct for nucleus rotation.

The time-averaged MSD of fibrillarin was calculated in MATLAB (@msdanalyzer^84^) by

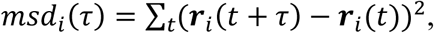

where ***r*** is the position vector at each time point and *τ* is the measurement time interval. As the MSD curves were not linear, indicating that the movement of fibrillarin was not Brownian motion, only 50% of the curves were taken to fit log(MSD) versus log(t), where we obtained the diffusion exponent α and diffusion coefficient *D* using R^2^ > 0.8 as the threshold. Additionally, the approximate time to diffuse 1 µm^2^ was calculated as

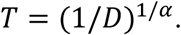

### Chromatin-modifying enzyme activity assays

Nuclear protein extractions were isolated from 500,000 fibroblasts seeded on elastic and viscoelastic substrates for 48 h using a nuclear extraction kit (EpiGentek, OP-0002) according to the manufacturer’s instructions. HAT, HDAC, H3K4 HMT, and H3K4 HDM activity were measured using the HAT activity/inhibition assay (EpiGentek, P-4003-048), HDAC activity/inhibition assay (EpiGentek, P-4034-096), HMT (H3K4 specific) activity/inhibition assay (EpiGentek, P-3002-1), and HDM (H3K4 specific) activity/inhibition assay (EpiGentek, P-3074-48), respectively. Per the manufacturer’s instructions, 20 µg of nuclear extract was added into the assay wells and incubated at 37 °C for 90-120 min. After adding the color developer solution, the absorbance was measured using a plate reader (Infinite 200Pro) at 450 nm for all the assays with the exception of the HDM (H3K4 specific) activity/inhibition assay where the fluorescence was measured at 530 EX/590 EM nm.

### Assay for transposase-accessible chromatin using sequencing (ATAC-seq)

#### Sample collection, preprocessing and sequencing

1,000,000 fibroblasts seeded on elastic or viscoelastic substrates were collected after 48 h and stored at −80 °C prior to sample processing, as previously described^34^. Frozen cells were thawed and washed once with PBS and then resuspended in 500 µL of cold PBS. The cell number was assessed by Cellometer Auto 2000 (Nexcelom Bioscience, Massachusetts, USA). 100,000 cells were then added to ATAC lysis buffer and centrifuged at 500 × g in a pre-chilled centrifuge for 5 min. Supernatant was removed and the nuclei were resuspended in 50 µL of tagmentation reaction mix by pipetting up and down. The reactions were incubated at 37 °C for 30 min in a thermomixer with shaking at 1000 rpm, and then cleaned up using the MiniElute reaction clean up kit (Qiagen). Tagmented DNA was amplified with barcoded primers. Library quality and quantity were assessed with Qubit 2.0 DNA HS Assay (Thermo Fisher), Tapestation High Sensitivity D1000 Assay (Agilent Technologies), and QuantStudio 5 System (Applied Biosystems). Equimolar pooling of libraries was performed based on QC values and sequenced on an Illumina NovaSeq (Illumina, California, USA) with a read length configuration of 150 PE for [100]M PE reads (50 m in each direction) per sample.

#### Data preprocessing, mapping, peak calling and downstream analysis

FASTQ files were trimmed with Trimmomatic^78^ after quality control checking. Pair-ended reads were then aligned to mouse reference genome (mm10) with Bowtie2^85^. Mitochondrial reads and PCR duplicates were removed using SAMtools^86^ and Picard (http://broadinstitute.github.io/picard/) respectively. Peaks were called using MACS3^87^ (q < 0.001), and only peaks outside the ENCODE blacklist regions were kept. All peaks from all samples were combined and merged using BEDtools. FeatureCounts^80^ was used to count the mapped reads for each sample. Peaks that were up- or downregulated in different conditions were defined using DESeq2^81^ with P_adj_ = 0.05 as the threshold and annotated using nearest genes and genomic features. Gene ontology (GO) enrichment analysis was then performed toward genes related to up- and down-regulated peaks using the enrichGO function in clusterProfiler package^82^. Peaks located at *cis*-regulatory elements related to genes of interest (± 5 kb region) were visualized using Integrative Genomics Viewer (IGV)^88^ to demonstrate up- or down-regulated differential peaks. To compare the ATAC signals in Ascl1-, MyoD-, ETV2-, and Oct4-target binding sites across the genome in elastic and viscoelastic group, publicly available datasets (GSE43916, GSE157339, GSE168521, and GSE85602) were used to obtain the shared regions between the recognized peaks in these datasets and the differentially expressed ATAC peaks.

### Chemical inhibition of histone deacetylase (HDAC)

To elucidate the role of HDAC in viscoelasticity-mediated iN reprogramming, cells cultured on tissue culture treated (TC) plates or gels for one day were treated with valproic acid (Cayman Chemical, 2.5 mg ml^−1^) for 24 h. To examine the inhibition effects, after 24 h, cells on TC plates were collected for Western blotting analysis of AcH3 level. For reprogramming experiments, after 24 h treatment with valproic acid, medium was changed to MEF medium with Dox (2 µg ml^−1^, from Sigma) to induce transgene expression (marked as day 0). After 24 h, the medium was changed to N3 medium and cells were cultured until day 7, at which point samples were fixed and immunostained for Tubb3.

### Small interfering RNA (siRNA) knockdown of lamin A

For lamin A siRNA knockdown, cells were plated on TC plates or elastic and viscoelastic substrates for 24 hours before adding siRNA. RNA interference was performed using ON-TARGETplus siLMNA (Dharmacon, L-040758-00-0005), and transfections were carried out using Lipofectamine 3000 reagent (Thermo Fisher, L3000015) according to the manufacturer’s protocol. Briefly, for 6 wells in a 24-well plate, 500 μl Opti-MEM medium (Thermo Fisher, 31985062) was mixed with 10 μl Lipofectamine 3000 reagent and incubated at 37 °C for 5 minutes. 10 μg siRNA was then added to the mixture and incubated at 37 °C for 20 minutes. The DNA-lipid complexes were added to 2.7 ml DMEM without FBS and penicillin/streptomycin, added to the cells, and incubated at 37 °C for 12 hours. The media were then replaced with MEF medium and cultured for 12 h. To examine the efficiency of the knockdown, cells on TC plates were collected for Western blotting analysis of lamin A/C expression. For reprogramming experiments, 12 h after removing medium containing the plasmid, medium was changed to MEF medium with Dox (2 µg ml^−1^, from Sigma) to induce transgene expression (marked as day 0). After 24 h, the medium was then changed to N3 medium and cells were cultured until day 7 to determine the reprogrammin efficiency.

### Quantitative reverse transcriptase-polymerase chain reaction

RNA was isolated from samples using Trizol (Ambion) according to the manufacturer’s instructions. For cDNA synthesis, 500 ng of RNA was reverse transcribed using Maxima First Strand cDNA Synthesis Kit (Thermo Fisher Scientific). Template DNA was amplified using Maxima SYBR Green/Fluorescein qPCR Master Mix (Thermo Fisher Scientific) on a CFX qPCR machine (Bio-Rad). qRT-PCR data were analyzed using CFX Manager 3.1 (Bio-Rad) and gene expression levels (endogenous *Ascl1*: forward primer, CAACCGGGTCAAGTTGGTCA; reverse primer, CTCATCTTCTTGTTGGCCGC; *Tubb3*: forward primer, GCCGCTAGAGGTGAAATTC TTG; reverse primer, CATTCTTGGCAAATGCTTTCG) were normalized to the expression level of *18S* (forward primer, GCCGCTAGAGGTGAAATTCTTG; reverse primer, CATTCTTGGCAA ATGCTTTCG).

### Western blotting analysis

Alginate lyase (Sigma) was dissolved in PBS to make a 15 U ml^−1^ solution. The lyase solution was cooled down and added to the gels after washing gels with cold PBS once to dissolve the gels and collect the cells. Cells were then lysed in RIPA buffer (Thermo Scientific, 89900) along with protease & phosphatase inhibitor cocktail (Thermo Scientific, 78440) on ice. Protein lysates were centrifuged to pellet cell debris, and the supernatant was collected and used for further analysis. Protein samples were run using SDS-PAGE and transferred to polyvinylidene fluoride membranes. Membranes were blocked in 5% nonfat milk and incubated with primary antibodies (**Table S1, Supplementary Information**) overnight on ice. Membranes were washed with Tris-Buffered Saline + 0.05% Tween-20 and incubated with HRP-conjugated IgG secondary antibodies (Santa Cruz Biotechnologies) for 1 h. Protein bands were visualized using Western Lightning Plus-Enhanced Chemiluminscence Substrate (Perkin Elmer Life & Analytical Sciences) and imaged on a ChemiDoc XRS system (Bio-Rad).

### Statistical analysis

The data were presented as mean plus or minus one standard deviation, where n ≥ 3. The data corresponding to the cell spreading area, nuclear volume, chromatin compaction index, and histone quantification experiments were displayed as truncated violin plots to demonstrate the distribution of data. The violins were drawn with the ends at the quartiles and the median as a horizontal line in the violin. Comparisons among values for groups greater than two were performed using a one-way or two-way analysis of variance (ANOVA) and differences between groups were determined using the following multiple comparison tests: Dunnett’s, Tukey’s, and Sidak’s post-hoc test. For comparison between two groups, a two-tailed, unpaired t-test was used. For all cases, p-values less than 0.05 were considered statistically significant. GraphPad Prism 9.0 were used for all statistical analysis.

## Data Availability

Sequencing data will be available at the GEO after publication. Previously published data that were used in this paper are available at the GEO under accession codes GSE43916, GSE157339, GSE168521, and GSE85602. Source data are provided with this paper. All other data supporting the findings of this study are available from the corresponding author upon reasonable request.

## Code availability

Codes utilized for image analysis and sequencing data analysis are available on the lab website (https://li-lab.seas.ucla.edu/requestform/).

## Acknowledgements

This work was supported, in part, by a UCLA Eli and Edythe Broad Center of Regenerative Medicine and Stem cell Research Innovation Award and grants from the National Institute of Health (GM143485 and NS130677 to S.L.), the National Science Foundation (BRITE Fellow Award CMMI-2135747 to A.R.), and the Department of Defense (Ovarian Cancer Research Fund TEAL Expansion Award to A.R.). The authors acknowledge the support by the NIH/NCI under award number P30CA016042 and the Advanced light Microscopy and Spectroscopy Laboratory at the California NanoSystems Institute. The content is solely the responsibility of the authors and does not necessarily represent the official views of the NIH.

## Author Contributions

Y.W. and S.L. conceived the study. Y.W., Y.S., J.S., D.Q., A.R. and S.L. designed the experiments. Y.W., Y.S., J.S., T.H., X.L., A.Z., S.C., R.M., D.Q., X.H., and K.Y. performed the experiments. Y.W., Y.S., J.S., T.H., and X.L. analyzed the data. Z.F., J.E. and L.G. provided the reagents. Y.W., Y.S., J.S., T.H., D.Q., Z.F., J.E., L.G., A.R., Z.G., and S.L. contributed to the data interpretation and discussion. Y.W., J.S., and S.L. wrote the manuscript.

