## Supplementary Information for "Viscoelastic Extracellular Matrix Enhances Epigenetic Remodeling and Cellular Plasticity"

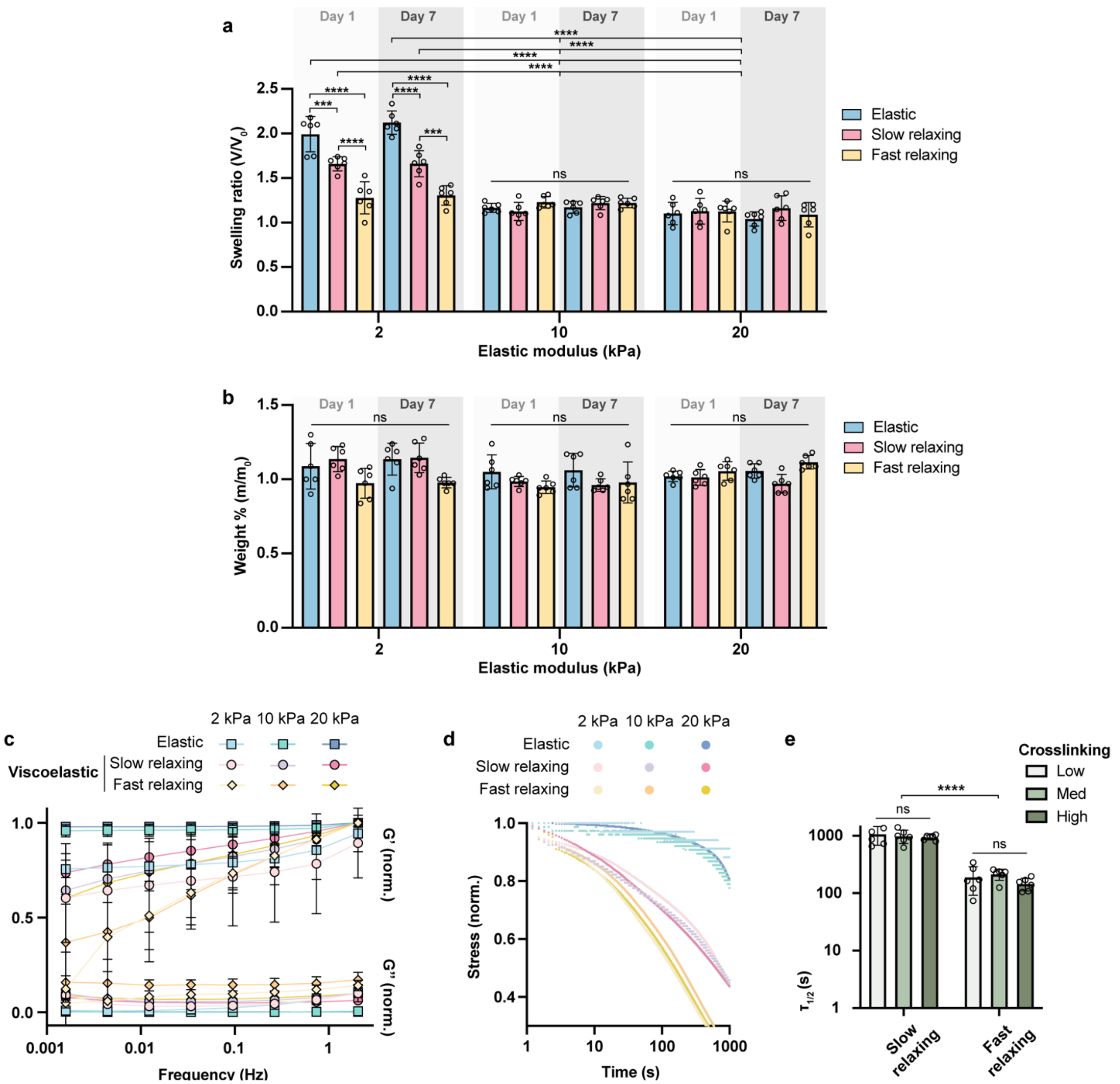


**Supplementary Fig. 1 | Charaterization of algiante-based gel substrates. a**, Swelling of gels with different type of crosslinking after 1 and 7 days of DMEM immersion, quantified by the ratio of the volume of gels at day 1 or day 7 to their volumes before immersion (n = 6 batches of gels). **b**, Weight ratio of the dry mass of gels after 1 and 7 days of DMEM immersion to the dry mass of gels before immersion (n = 6 batches of gels). **c**, Rheology measurements of the storage modulus and loss modulus of covalently (elastic) and ionically (viscoelastic, slow relaxing and fast relaxing) crosslinked alginate hydrogels with varying stiffnesses (n = 3 batches of gels). **d**, Stress of various gels changing with time under a constant strain. **e**, The time taken to relax the initial stress to half of the original value ($\tau_{1/2}$ for viscoelastic gels of different degree of crosslinking, as determined by compression testing (n = 6 batches of gels). In **a-c** and **e**, data are shown as mean ± s.d. Two-way ANOVA using Tukey's correction for multiple comparisons was used to determine the significance between the groups (***$P < 0.001$, ****$P < 0.0001$).

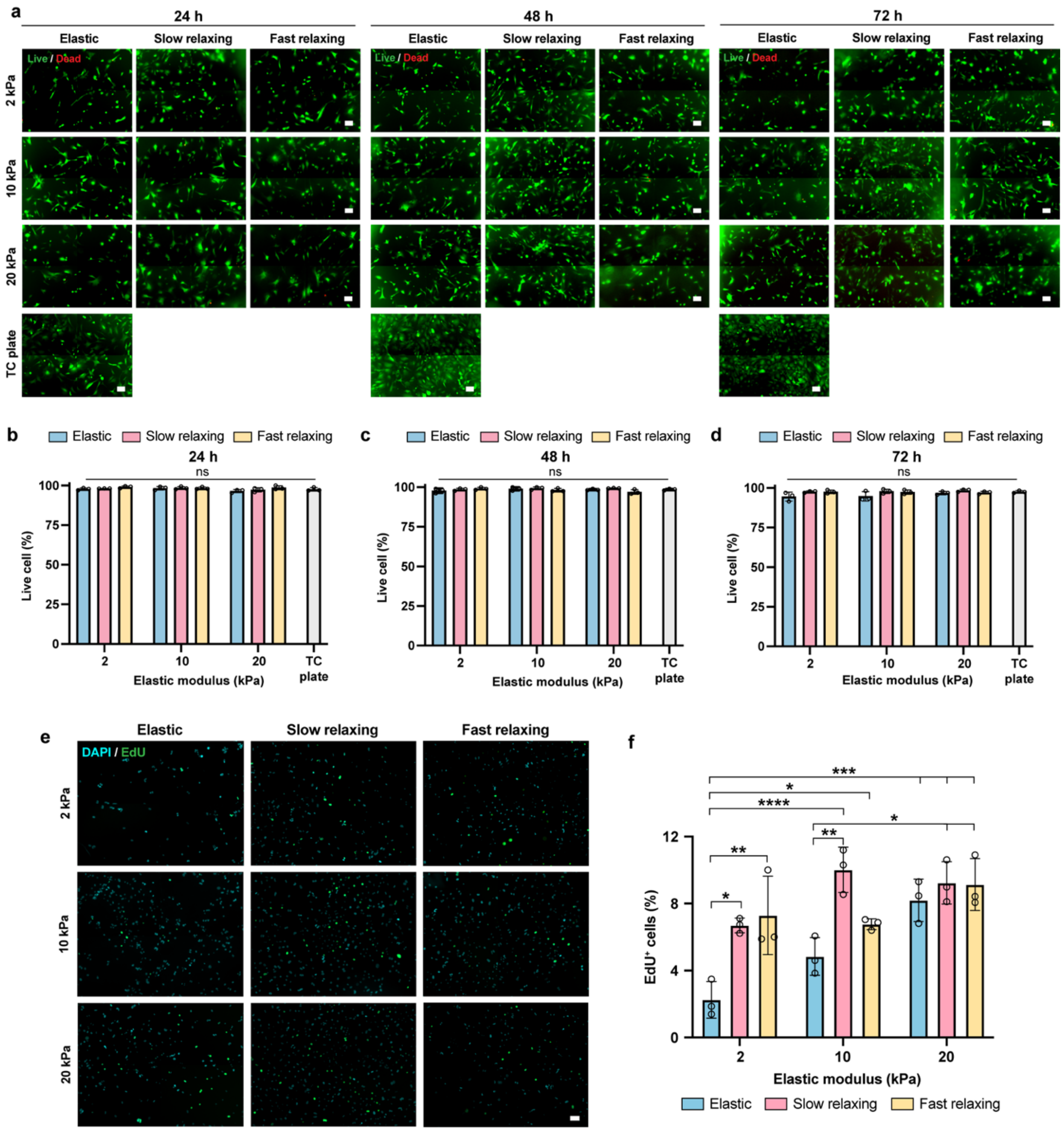


**Supplementary Fig. 2 | Viability and proliferation of fibroblasts on various substrates. a**, Live/dead staining showing cell viability at 24, 48, and 72 hours after cell seeding (left to right). Scale bar, 100 $\mu$m. **b-d**, Quantification of viability of fibroblasts cultured on gel substrates and tissue culture (TC) plates for 24 (**b**), 48 (**c**), and 72 (**d**) hours. n = 3 independent experiments. **e**, EdU staining (**e**) of fibroblasts cultured on various substrates at 48 hours. Scale bar, 100 $\mu$m. **f**, Quantification of the percentage of EdU$^+$ cells (n = 3 independent experiments). Two-way ANOVA using Tukey's correction for multiple comparisons was used to determine the significance between the groups (*$P < 0.05$, **$P < 0.01$, ***$P < 0.001$, ****$P < 0.0001$).

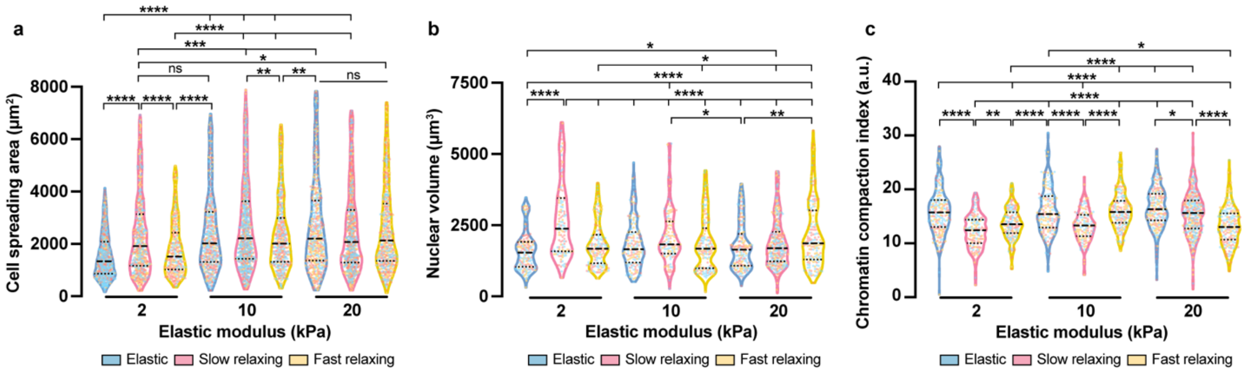


**Supplementary Fig. 3 | Substrate viscoelasticity regulates cell spreading, nuclear volume, and chromatin organization. a**, Cell spreading area of fibroblasts cultured on various substrates for 24 hours (n= 810, 676, 381, 608, 893, 489, 589, 831, 604 cells per condition from 3 independent experiments). **b**, Nuclear volume of fibroblasts cultured on gel substrates for 24 hours (n = 187, 140, 167, 154, 136, 173, 193, 346, 174 cells per condition from 3 independent experiments). **c**, Chromatin compaction index of fibroblasts on substrates at 24 hours, quantified by the ratio of integrated fluorescence intensity of DAPI staining to nuclear volume (n = 186, 140, 163, 152, 133, 172, 193, 344, 174 cells per condition from 3 independent experiments). Truncated violin plots were used to demonstrate the distribution of data. The violins are drawn with the ends at the quartiles and the median as a horizontal line in the violin, where the curve of the violin extends to the minimum and maximum values in the data set. The color of dots inside the violins reflects the experimental replicates. Significance was determined by one-way ANOVA using Tukey's correction for multiple comparisons (**P* < 0.05, ***P*<0.01, ****P* <0.001, *****P* < 0.0001).

DAPI, orthogonal View

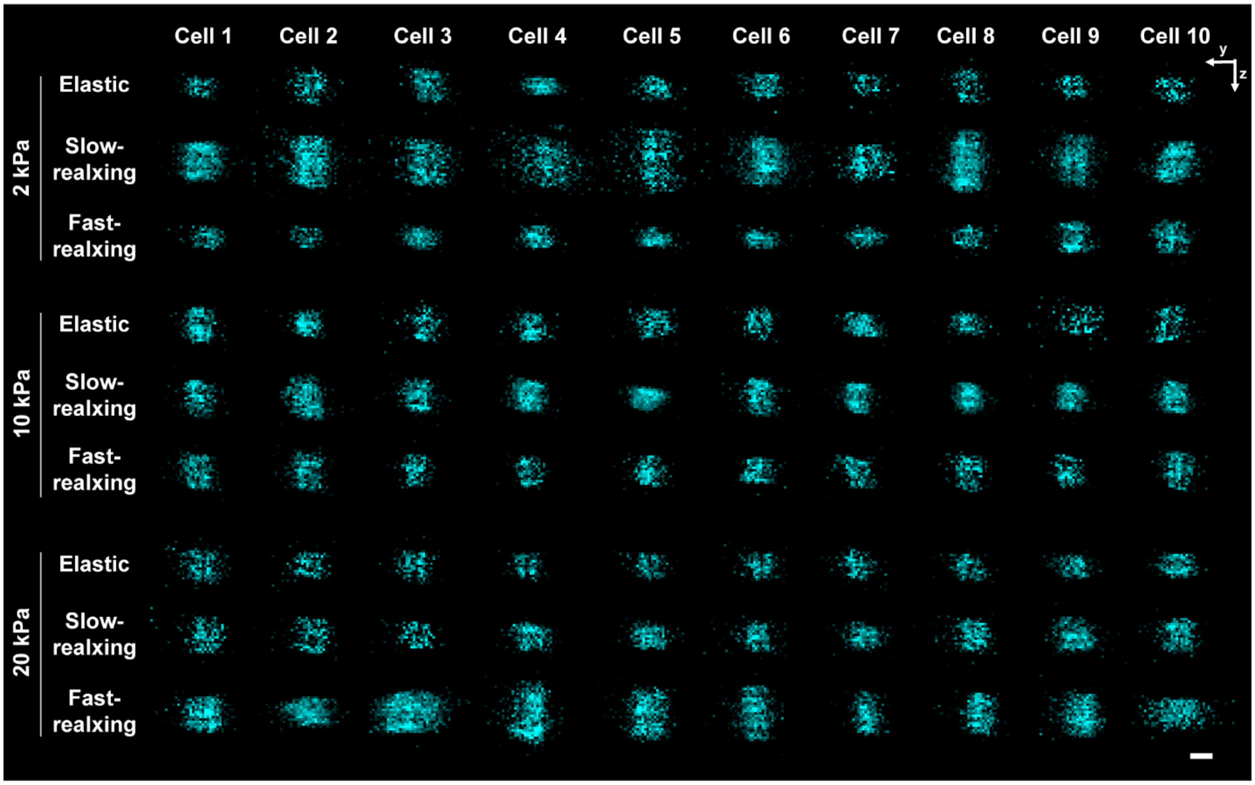


**Supplementary Fig. 4 | Orthogonal views of representative cell nuclei on various substrates.** Images were utilized to determine the nuclear volume of cells across various substrates. Scale bar, 10 $\mu$m.

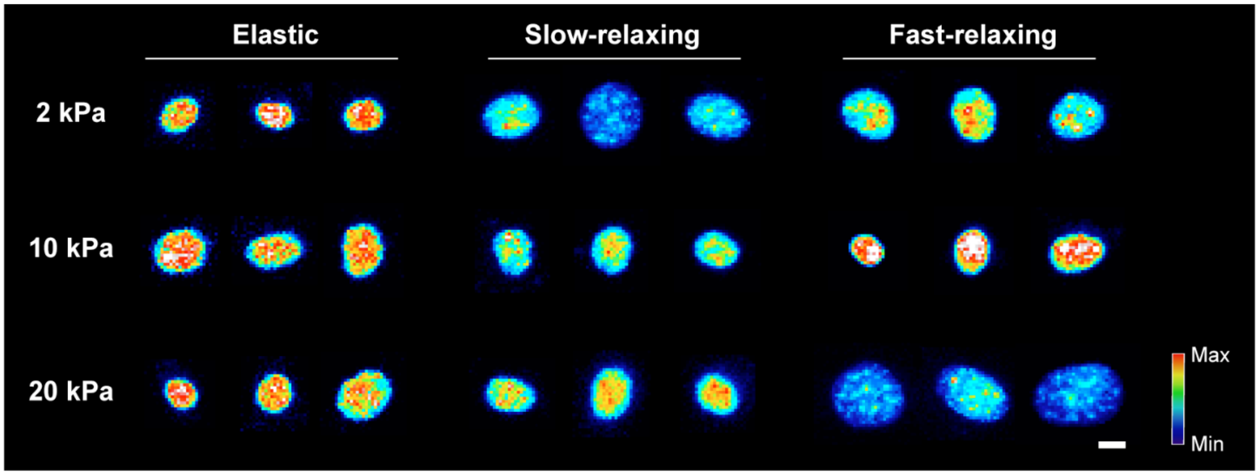


**Supplementary Fig. 5 | Representative images of DAPI-stained cell nuclei on various substrates.** Images were utilized to determine the chromatin compaction index of cells across various substrates. Scale bar, 10 $\mu$m.

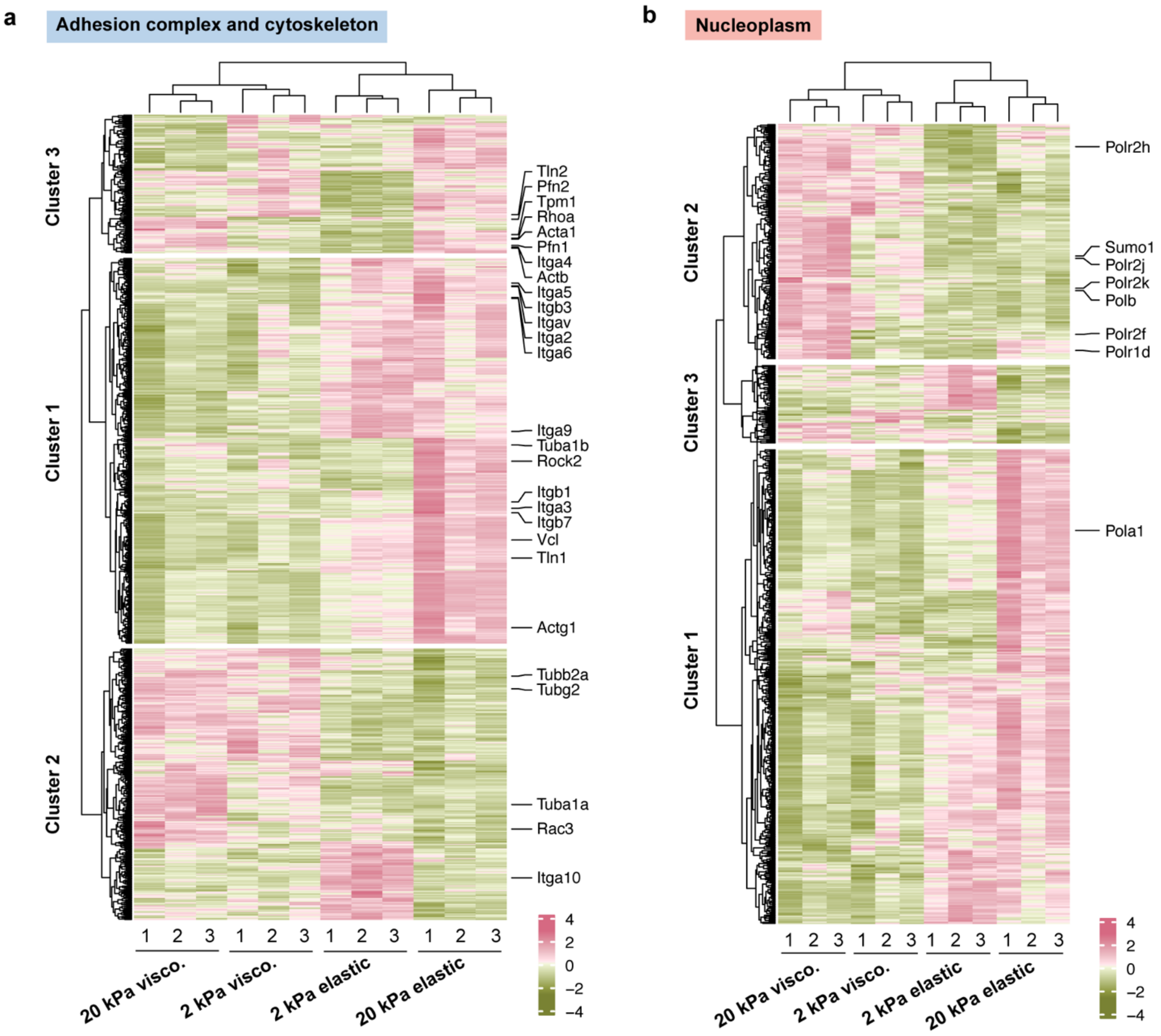


**Supplementary Fig. 6 | Differentially expressed genes (DEGs) in Adhesion complex and cytoskeleton and Nucleoplasm categories. a**, Heatmap of DEGs in the Adhesion complex and cytoskeleton category. **b**, Heatmap of DEGs in the Nucleoplasm category.

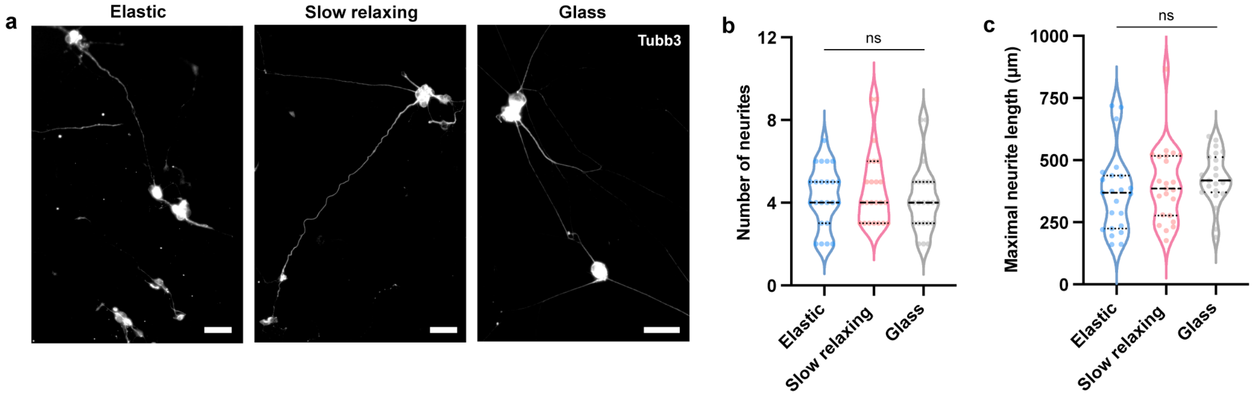


**Supplementary Fig. 7 | Morphological assessment of neurons derived on various substrates. a**, Tubb3-immunostained induced neuronal cells derived on elastic, slow-relaxing, and glass substrates after 3-4 weeks. Scale bar, 50 $\mu$m. **b-c**, Quantification of the number of neurites (**b**) and the maximal neurite length (**c**) of neuronal cells (n = 23 cells). In **b-c**, the violins are drawn with the ends at the quartiles and the median as a horizontal line in the violin, where the curve of the violin extends to the minimum and maximum values in the data set. Significance was determined by one-way ANOVA using Tukey's correction for multiple comparisons.

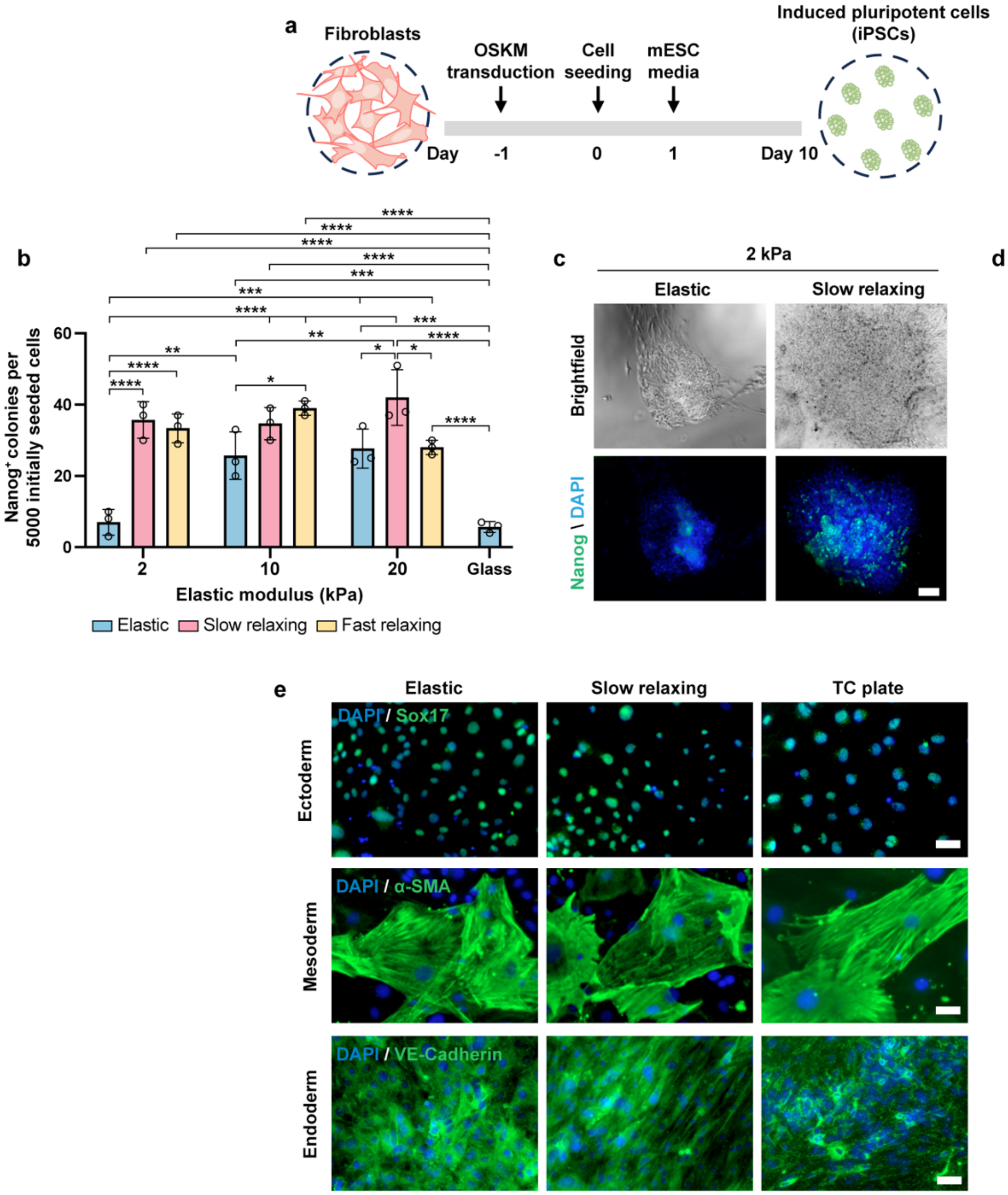


**Supplementary Fig. 8 | Substrate viscoelasticity promotes the reprogramming of fibroblasts into induced pluripotent stem cells (iPSCs). a**, Schematic illustration of the reprogramming timeline. **b**, Quantification of Nanog$^+$ colonies on elastic or viscoelastic substrates at day 10 (n = 3 independent experiments). **c**, Representative brightfield (top) and fluorescent (bottom) images of Nanog$^+$ colonies on 2-kPa gels at day 10. Scale bar, 100 $\mu$m. **d**, PCR analysis of *Nanog* mRNA expression in cells on various substrates at day 10 (n = 3 independent experiments). **e**, Fluorescent images of differentiated iPSCs that were immunostained for a neuronal lineage marker Sox17 (ectoderm, top), muscle lineage marker α-SMA (alpha smooth muscle actin, mesoderm, middle), and endothelial lineage marker VE-Cadherin (endoderm, bottom). Scale bar, 50 $\mu$m. In **b** and **d**, bar graphs show the mean ± s.d. In **b**, a two-way ANOVA using Tukey's correction for multiple comparisons was used to determine the significance between the groups. In **d**, a two-tailed, unpaired t test was used to determine statistical significance (*$P$ < 0.05, **$P$ < 0.01, ***$P$ < 0.001, ****$P$ < 0.0001).

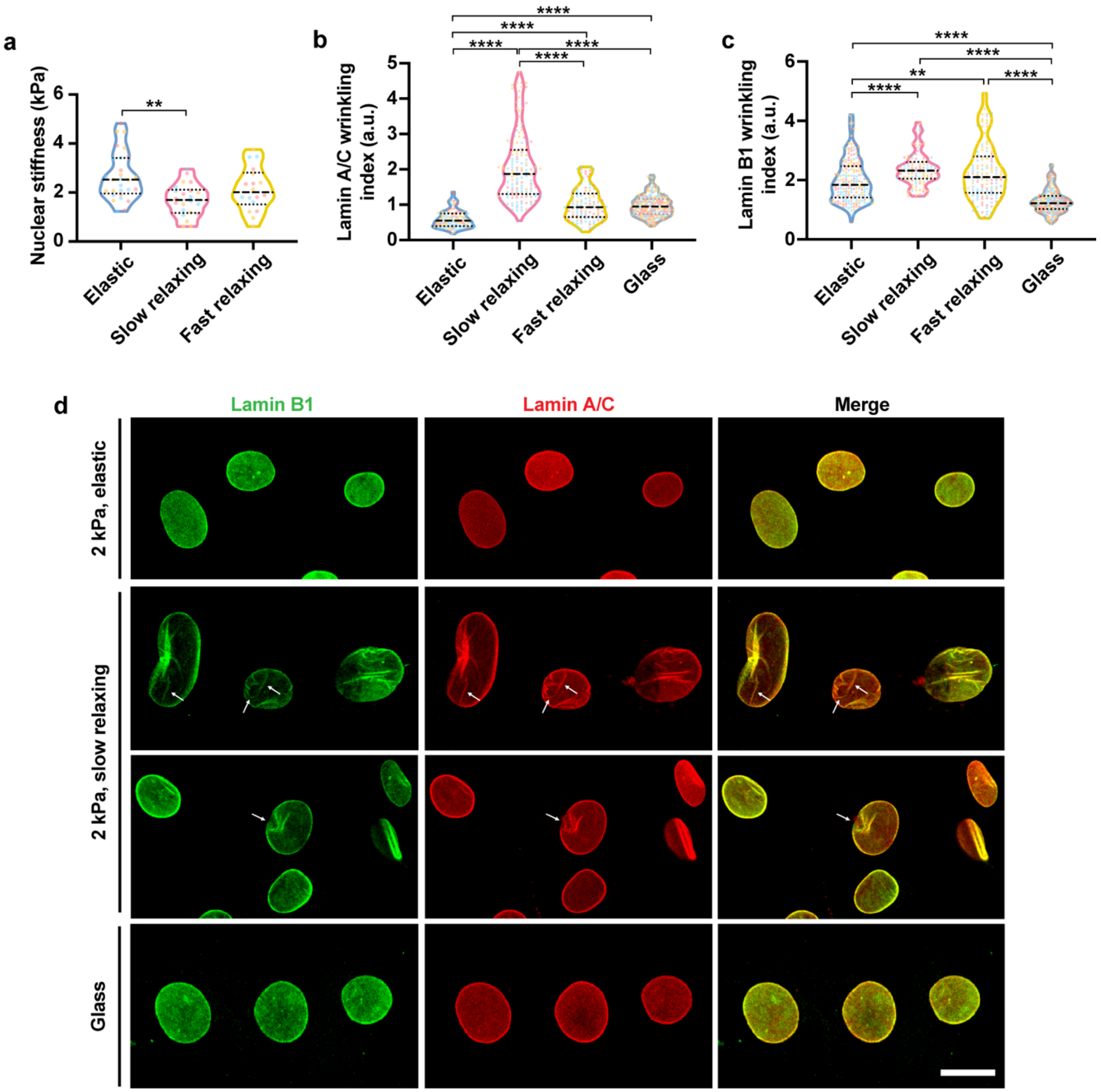


**Supplementary Fig. 9 | Substrate viscoelasticity remodels the nuclear lamina. a**, Nuclear stiffness of fibroblasts on 2-kPa substrates, as measured by nano-indentation (n = 29, 24, 18 cells per condition from 3 independent experiments). **b-c**, Quantification of nuclear wrinkling index on 2-kPa substrates indicated by lamin A/C immunostaining (**b**) or lamin B1 staining (**c**). n = 146, 145, 116 cells per condition from 3 independent experiments for lamin A/C; n = 197, 108, 147 cells per condition from 3 independent experiments for lamin B1. **d**, Co-staining of lamin A/C and lamin B1 on 2-kPa substrates and glass demonstrated the heterogeneity of the nuclear lamina. Scale bar, 20 $\mu$m. In **a-c**, the violins are drawn with the ends at the quartiles and the median as a horizontal line in the violin, where the curve of the violin extends to the minimum and maximum values in the data set. The color of dots inside the violins reflects the experimental replicates. Significance was determined by one-way ANOVA using Tukey's correction for multiple comparisons (**$P$<0.01, ****$P$ < 0.0001).

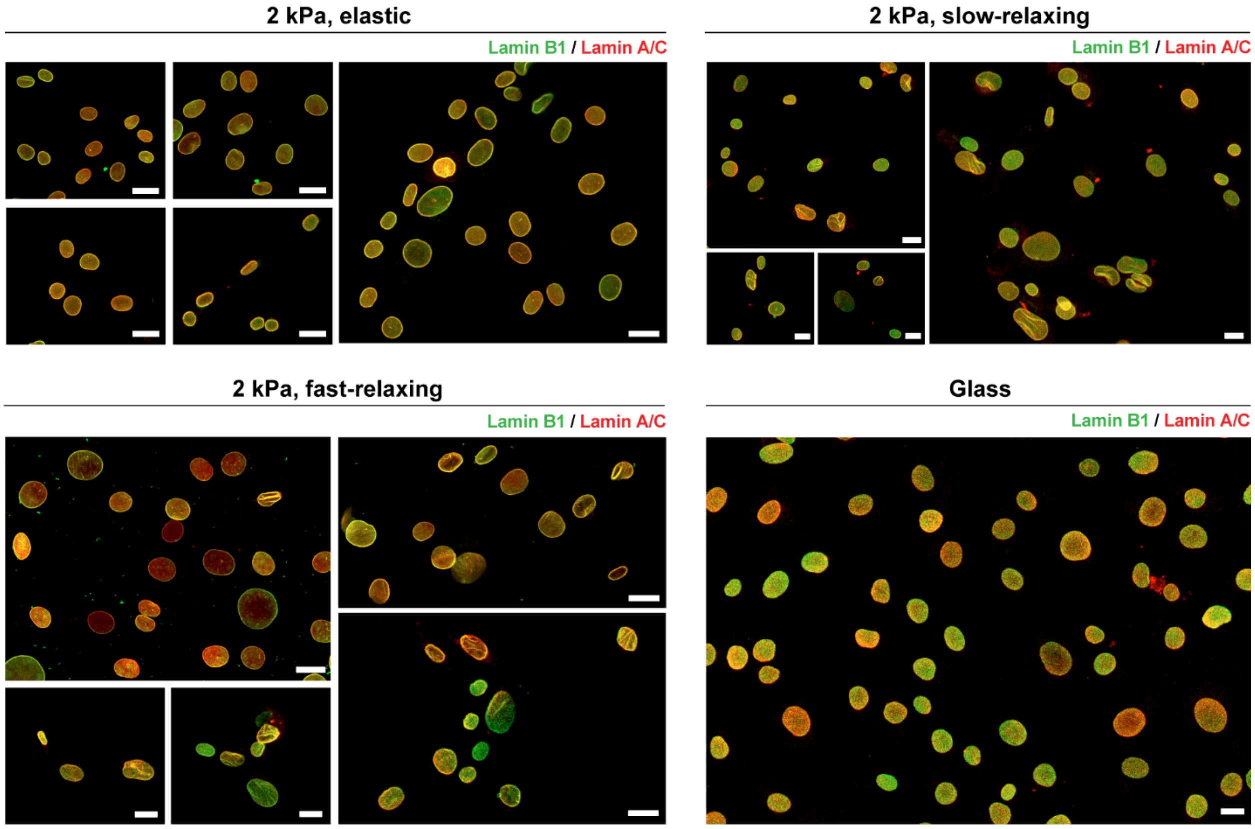


**Supplementary Fig. 10 | Additional representative images of fibroblasts co-stained for lamin A/C and lamin B1.** Cells were cultured on various substrates for 48 hours. Scale bar, 20 $\mu$m.

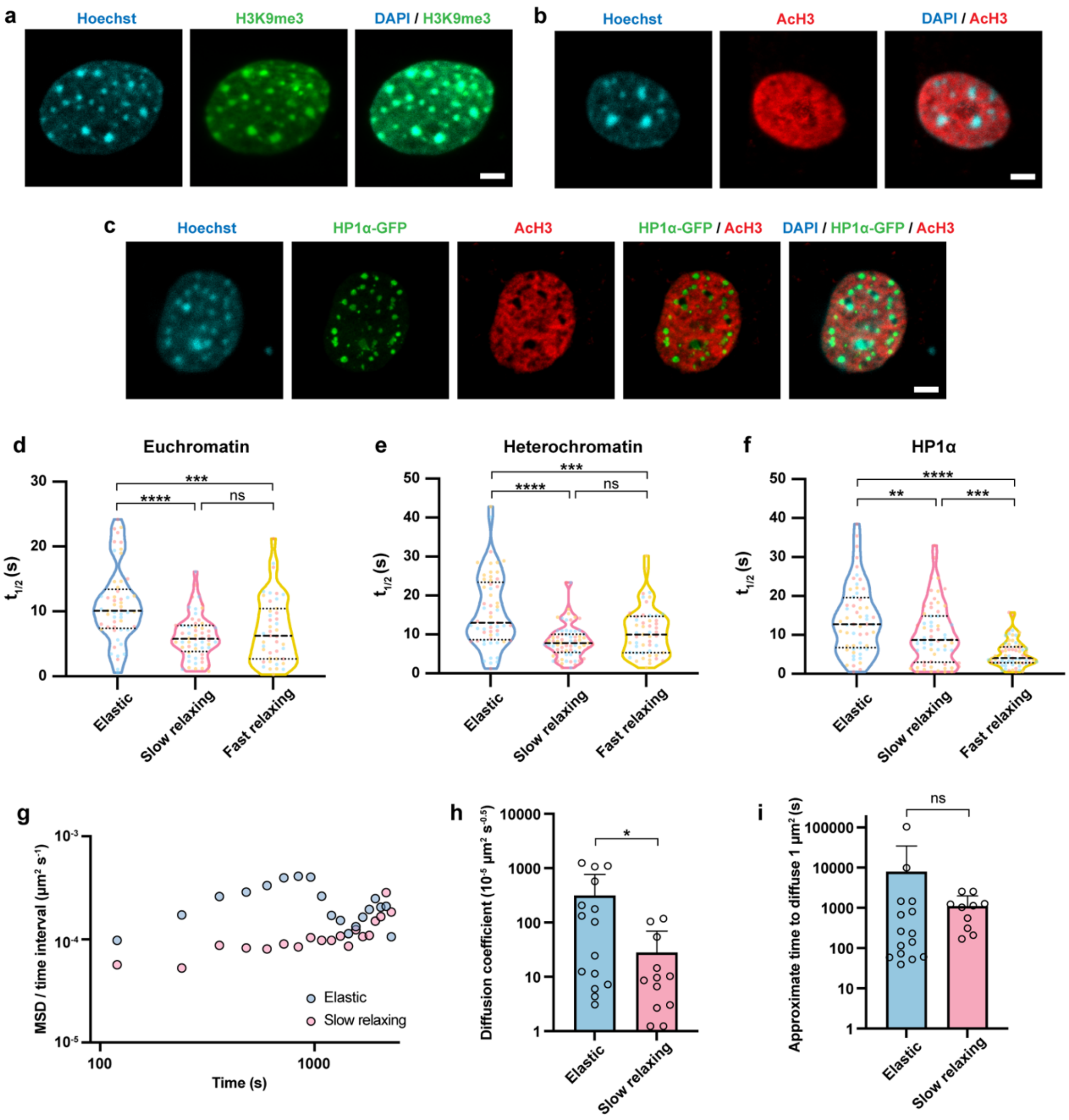


**Supplementary Fig. 11 | Substrate viscoelasticity enhances chromatin dynamics. a-b**, Immunostaining images of Hoechst-labeled cell nuclei with heterochromatic mark tri-methylated histone H3 on lysine 9 (H3K9me3, **a**) or euchromaic mark histone H3 acetylation (AcH3, **b**). Scale bar, 5 $\mu$m. **c**, Fluorescent images of the cell nucleus of fibroblasts expressing GFP-tagged HP1α (HP1α-GFP) labeled with Hoechst and immunostained for AcH3. Scale bar, 5 $\mu$m. **d-e**, Truncated violin plots showing the recovery half time of the photo-bleahced area in euchromatic (**d**) and heterochromatic regions (**e**) of fibroblasts on 2-kPa substrates. For euchromatic regions, n = 54, 56, 43 cells per condition from 3 independent experiments. For heterochromatic regions, n = 57, 72, 51 cells per condition from 3 independent experiments. **f**, Truncated violin plots showing the recovery half time of the GFP fluorescence in the cell nuclei of fibroblasts expressing HP1α-GFP (n = 56, 67, 68 cells per condition from 3 independent experiments). **g**, Mean square displasment (MSD) divided by time in fibroblasts expressing GFP-tagged fibrillarin as determined by using confocal microscopy to measure fibrillarin movement (n = 20, 19 cells per condition from 3 independent experiments). **h**, Diffusion coefficient of fibrillarin movement trajectories extracted from elastic and slow-relaxing substrates of 2 kPa (n = 15, 12 cells per condition from 3 independent experiments). Only the diffusion coefficients with fitting $R^2 > 0.8$ were kept for analysis. **i**, Approximate time to diffuse 1 $\mu$m calculated from fibrillarin movement trajectories on elastic and slow-relaxing gels of 2 kPa (n = 15, 10 cells per condition from 3 independent experiments). Only both the diffusion coefficients and diffusion exponents with fitting $R^2 > 0.8$ were kept for analysis. In **d-f**, truncated violin plots were used to demonstrate the distribution of data. The violins are drawn with the ends at the quartiles and the median as a horizontal line in the violin, where the curve of the violin extends to the minimum and maximum values in the data set. The color of dots inside the violins reflects the experimental replicates. In **d-f**, significance was determined by one-way ANOVA using Tukey's correction for multiple comparisons. In **h-i**, data are shown as mean $\pm$ s.d. and a two-tailed, unpaired t test was used to determine the statistical significance (*$P < 0.05$, **$P<0.01$, ***$P < 0.001$,****$P < 0.0001$).

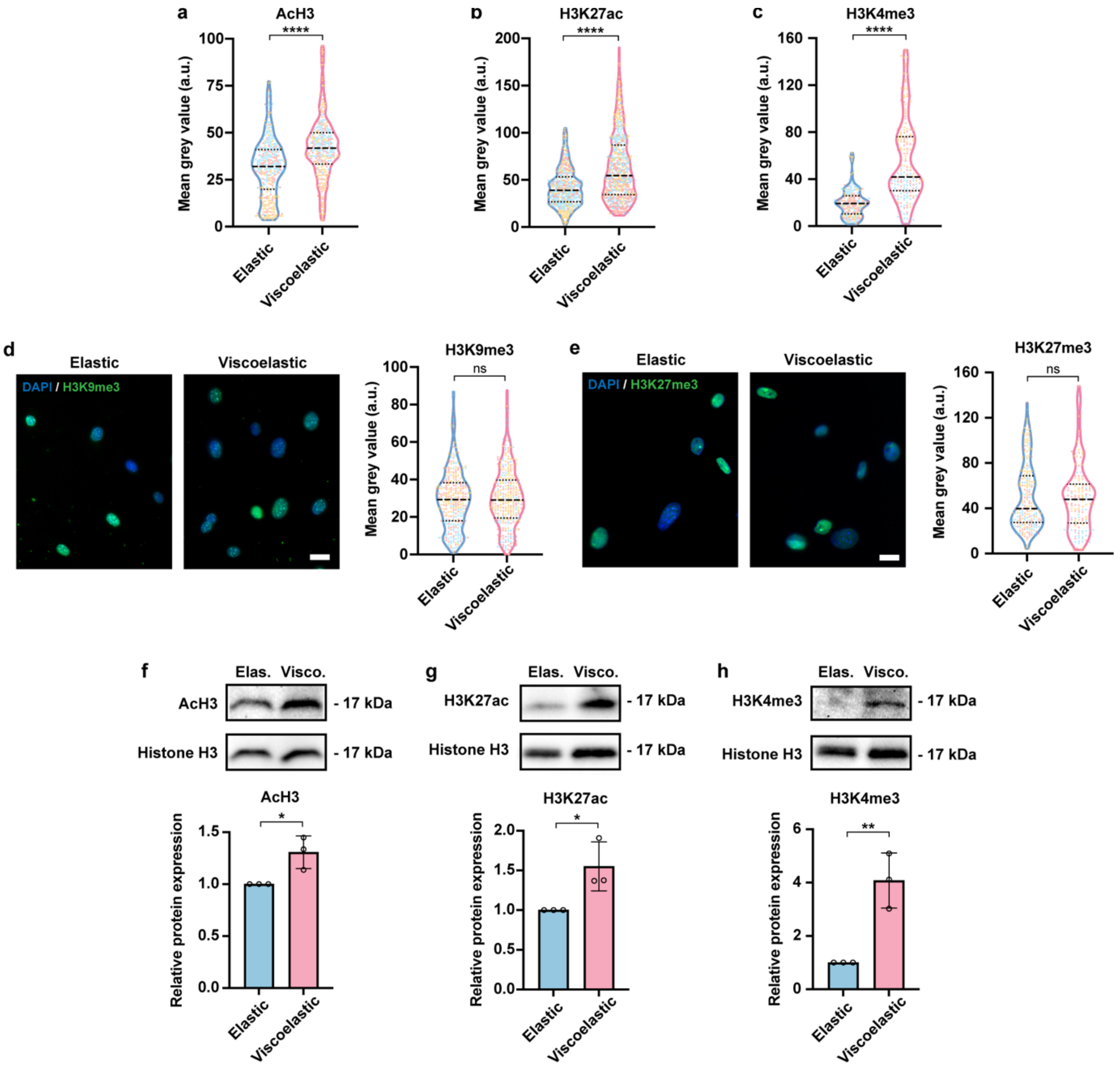


**Supplementary Fig. 12 | Substrate viscoelasticity modulates epigenetic marks. a-c**, Truncated violin plots showing the fluorescence intensity quantification of AcH3 (**a**), H3K27ac (**b**), and H3K4me3 (**c**) in mouse fibroblasts cultured on elastic or viscoelastic (slow relaxing) substrates of 2 kPa for 48 hours (n = 342, 302 cells from 3 independent experiments per condition for AcH3; 468, 626 cells from 3 independent experiments per condition for H3K27ac; 250, 180 cells from 3 independent experiments per condition for H3K4me3). **d-e**, Fluorescent images of H3K9me3 (**d**) and tri-methylated histone H3 on lysine 27 (H3K27me3, **e**) in fibroblasts on elastic or viscoelastic substrates of 2 kPa, with fluorescent intensity quantifications shown as truncated violin plots (n = 318, 301 cells from 3 independent experiments per condition for H3K9me3; 231, 142 cells from 3 independent experiments for H3K27me3). Scale bar, 20 $\mu$m. **f-h**, Western blotting analysis of AcH3 (**f**), H3K27ac (**g**), and H3K4me3 (**h**), where histone H3 served as loading control. Quantification of AcH3 (**f**), H3K27ac (**g**), and H3K4me3 (**h**) levels from blots, where the protein expression level was normalized to the corresponding histone H3 and elastic groups (n = 3 independent experiments). In **a-e**, truncated violin plots were used to demonstrate the distribution of data. The violins are drawn with the ends at the quartiles and the median as a horizontal line in the violin, where the curve of the violin extends to the minimum and maximum values in the data set. The color of dots inside the violins reflects the experimental replicates. In **f-h**, data are shown as mean ± s.d. and a two-tailed, unpaired t test was used to determine the statistical significance (**P* < 0.05, ***P*<0.01, ****P* < 0.001,*****P* < 0.0001).

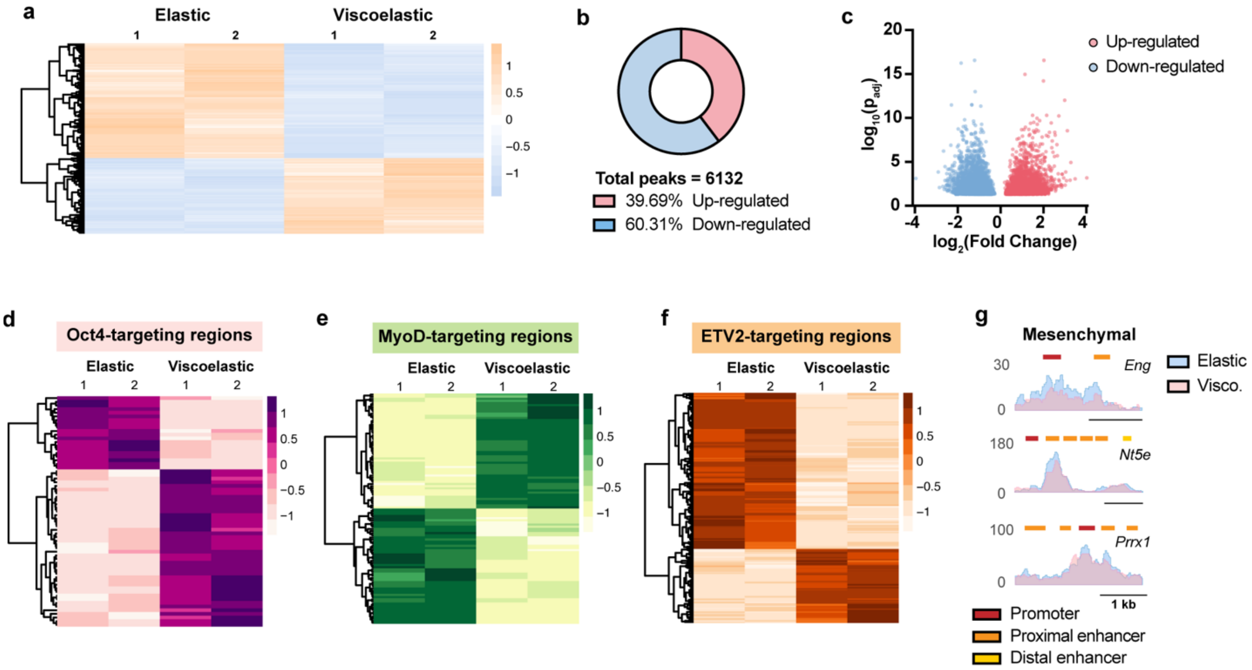


**Supplementary Fig. 13 | Substrate viscoelasticity regulates genome-wide chromatin accessibility towards specific lineages.** **a**, Heatmap of differentially expressed ATAC peaks from samples of fibroblasts cultured on elastic and viscoelastic substrates for 48 hours (n = 2 biological replicates). **b**, Percentage of up- and down-regulated peaks of viscoelastic groups in comparison with elastic groups among all the differentially expressed peaks. **c**, Volcano plots showing the fold changes and adjusted p-values of differentially expressed peaks. Each dot represents a peak. **d-f**, Heatmaps showing the differentially expressed ATAC signals in Oct4- (**d**), MyoD- (**e**), and ETV2- (**f**) targeting genomic sites. **g**, Peaks around the promoters of *Eng* (CD105), *Nt5e* (CD73), and *Prrx1* in fibroblasts cultured on elastic and viscoelastic substrates. Scale bar, 1 kb.

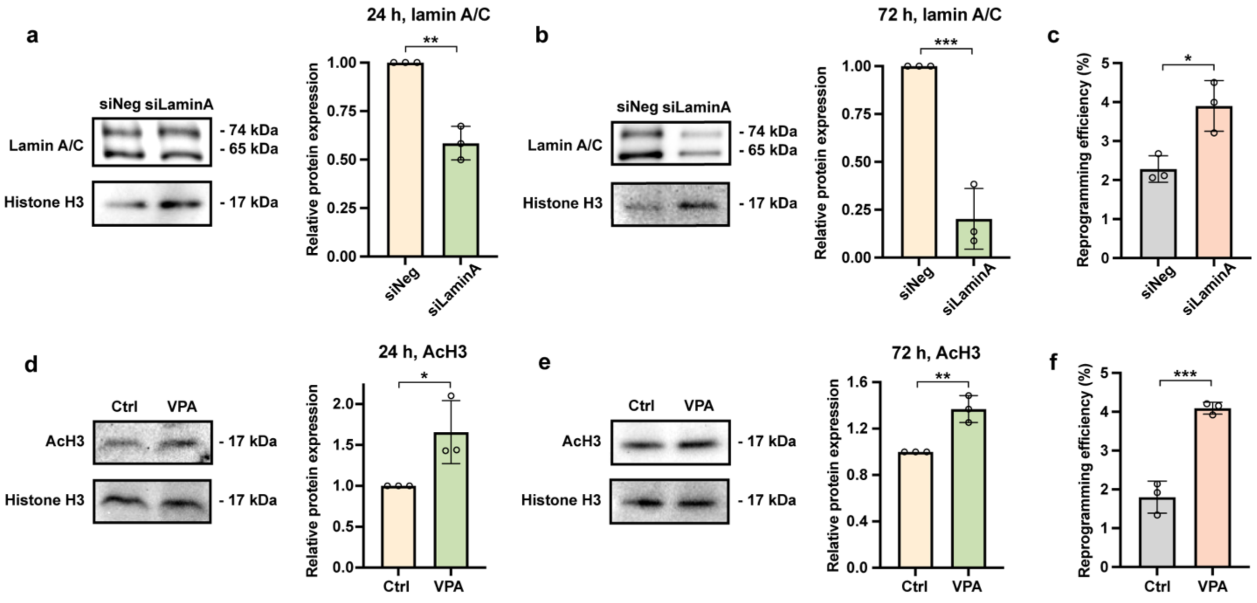


**Supplementary Fig. 14 | Silencing *Lmna* and inhibition of HDAC activity promote iN reprogramming. a-b**, Western blotting analysis of lamin A/C expression in fibroblasts transfected with a negative control or *Lmna* siRNA and cultured on TC plates for 24 hours (**a**) or 72 hours (**b**), where histone H3 served as a loading control. Quantification of lamin A/C expression from blots, where the protein expression level was normalized to histone H3 and ctrl groups, respectively (n = 3 independent experiments). **c**, Reprogramming efficiency of BAM-transduced fibroblasts transfected with a negative control or *Lmna* siRNA and cultured on glass for 7 days (n = 3 independent experiments). **d-e**, Western blotting analysis of AcH3 level in fibroblasts treated without or with 2.5 mg $ml^{-1}$ valproic acid (VPA, an HDAC inhibitor) and cultured on TC plates for 24 hours (**d**) or 72 hours (**e**). Inhibitors were removed after 24 hours of treatment. Quantification of AcH3 level from blots, where the protein expression level was normalized to histone H3 and ctrl groups, respectively (n = 3 independent experiments). **f**, Reprogramming efficiency of BAM-transduced fibroblasts cultured on glass in the absence or presence of VPA for 7 days, where VPA was removed after 24 hours of treatment (n = 3 independent experiments). Two-tailed, unpaired t tests were used to determine the statistical significance (*$P < 0.05$, **$P<0.01$, ***$P < 0.001$).

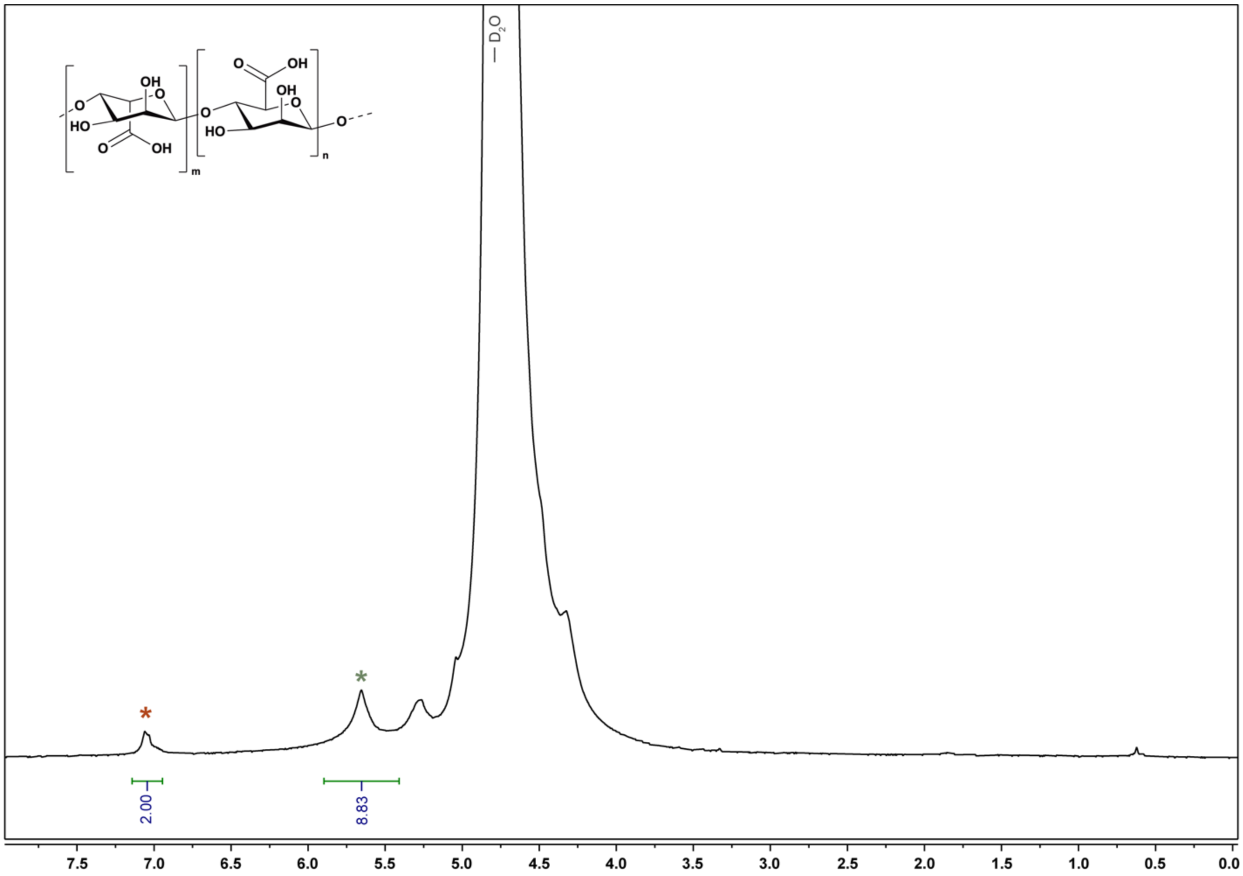


**Supplementary Fig. 15 | NMR spectra of unconjugated sodium alginate.** $^1$H NMR spectrum (383 K, 500 MHz, $D_2O$) of pristine alginate. Red star, maleic acid standard peak. Green star, G-1 alginate peak.

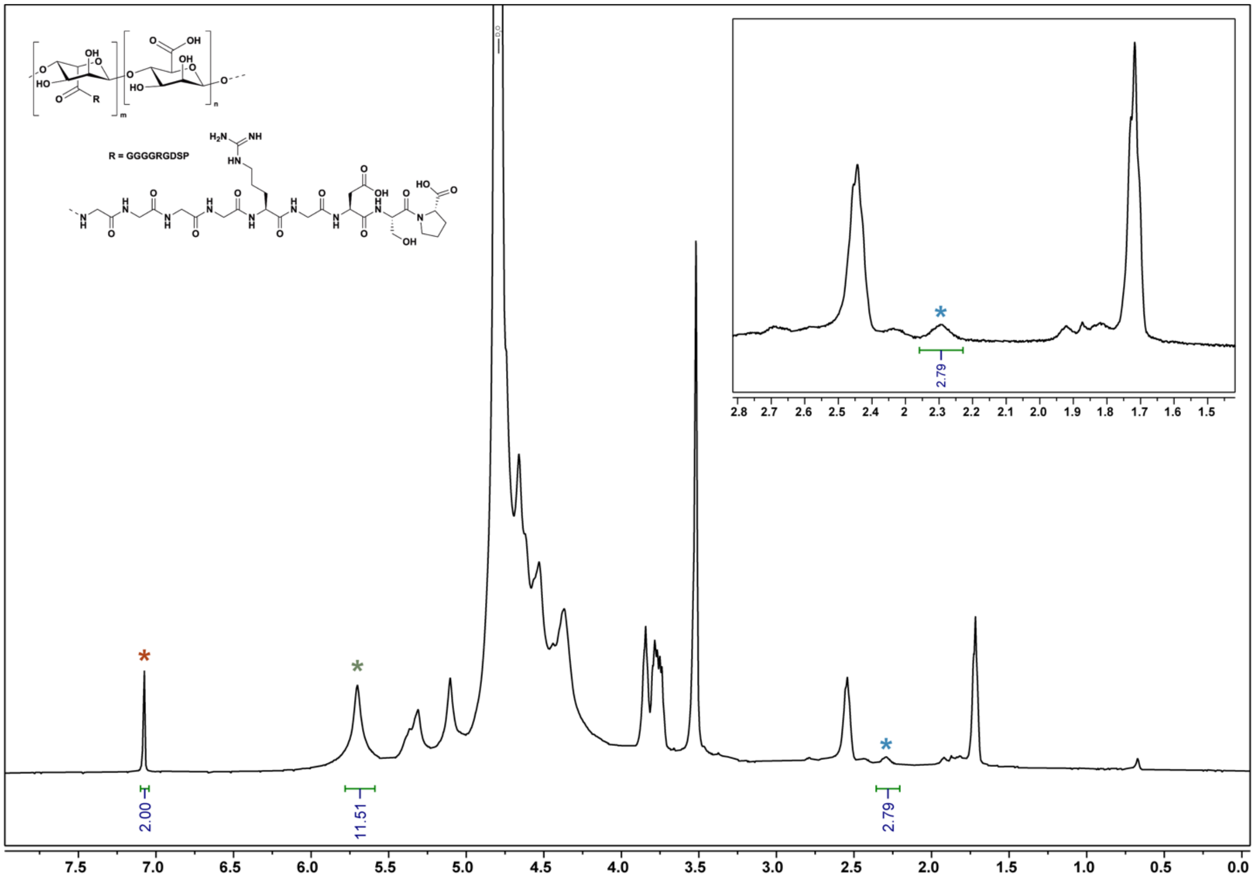


**Supplementary Fig. 16 | NMR spectra of RGD-coupled sodium alginate.** $^{1}$H NMR spectrum (383 K, 500 MHz, $D_2O$) of RGD-coupled alginate. Inset: peak at 2.30 ppm corresponding to the pyrrolidine side chain of proline. Red star, maleic acid standard peak. Green star, G-1 alginate peak. Blue star, proline peak.

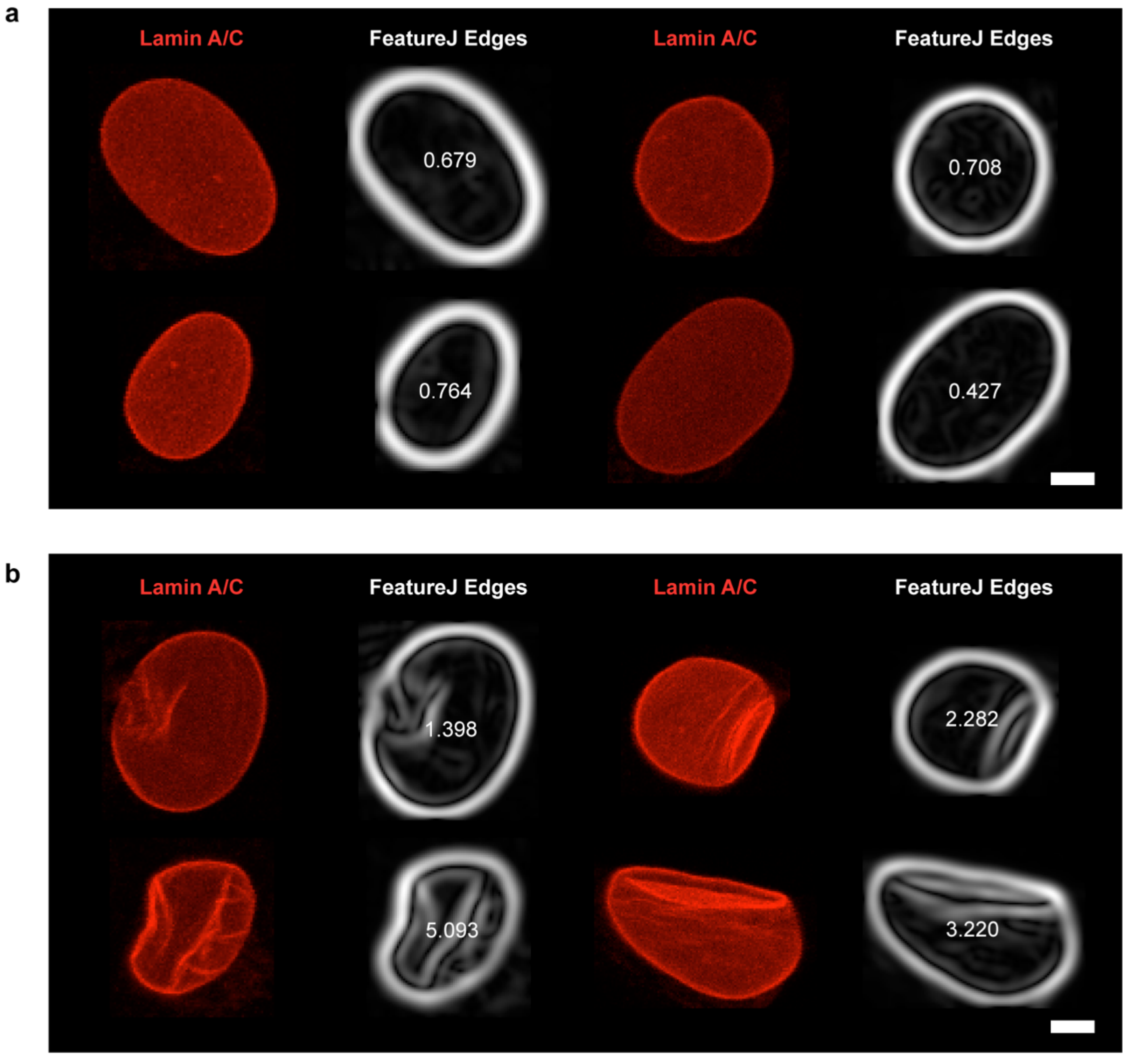


**Supplementary Fig. 17 | Quantification of lamin wrinkling using Fiji FeatureJ Edge detection.** Fluorescent images of lamin A/C staining of no-wrinkle cells (**a**) and wrinkled cells (**b**) and corresponding output images from FeatureJ Edge tracking. White numbers indicate the lamin wrinkling indices obtained from the measurement. Scale bar, 5 μm.

### Supplementary Note 1 – Average ligand density on alginate chains

As the density of cell-adhesive ligands can influence cellular behaviors in adherent cells, determining the ligand density in tissue-mimicking hydrogels is important to allow researchers to cross-compare the experimental findings. To calculate the average ligand density of polymer chains in our alginate-based hydrogel system, we performed quantitative nuclear magnetic resonance (qNMR) on unconjugated and RGD-coupled sodium alginate using maleic acid as the standard (**Supplementary Figs. 15-16**) and calculated the RGD peptide density as described previously[1].

The ratio of RGD peptide to maleic acid (NMR standard) was calculated as

$$concentration\ of\ peptide = \frac{m_S}{m_A} \cdot \frac{I_A}{I_S} \cdot \frac{1}{Mw_S} \cdot \frac{N_S}{N_A} \approx 8.017 \times 10^{-4}\ mol\ g^{-1},$$

where *m* is the mass, *I* is the integration of NMR peak, *Mw* is the molecular weight, and *N* is the proton number corresponding to the peak; *S* and *A* refer to standard and analyte respectively.

Similarly, the ratio of alginate to maleic acid was calculated as

$$concentration\ of\ alginate = \frac{m_S}{m_A} \cdot \frac{I_A}{I_S} \cdot \frac{1}{Mw_S} \cdot \frac{N_S}{N_A} \approx 1.295 \times 10^{-5}\ mol\ g^{-1}.$$

Therefore, the ligand density (number of RGD peptide per alginate chain) is

$$ligand\ density = \frac{concentration\ of\ peptide}{concentration\ of\ alginate} \approx 61.89.$$

**Table S1. Antibody Information**

| Antibody | Vendor | Catalog # | Dilution |
|---|---|---|---|
| Lamin A/C | Santa Cruz | sc-376248 | 1:100 (IF)<br>1:2000 (WB) |
| Lamin A/C | Thermo Scientific | PIMA535284 | 1:200 (IF) |
| Lamin B1 | Santa Cruz | sc-374015 | 1:150 (IF)<br>1:2000 (WB) |
| AcH3 | Millipore | 06-599 | 1:400 (IF)<br>1:5000 (WB) |
| H3K27ac | Abcam | ab4729 | 1:600 (IF)<br>1:5000 (WB) |
| H3K4me3 | Sigma | 07-473 | 1:400 (IF)<br>1:3000 (WB) |
| H3K9me3 | Abcam | ab8898 | 1:400 (IF) |
| H3K27me3 | Abcam | ab192985 | 1:400 (IF) |
| Tubb3 | Biolegend | 801202 | 1:1000 (IF) |
| MAP-2 | Sigma | M9942 | 1:200 (IF) |
| Synapsin | Abcam | ab64581 | 1:100 (IF) |
| vGLUT1 | Sigma | MAB5502 | 1:200 (IF) |
| GABA | Sigma | A2052 | 1:500 (IF) |
| Nanog | Abcam | ab214549 | 1:300 (IF) |
| $\alpha$-SMA | Abcam | ab7817 | 1:350 (IF) |
| Sox17 | Sigma | 09-038-I | 1:200 (IF) |
| VE-Cadherin | Invitrogen | 50-128-90 | 1:100 (IF) |
| Histone H3 | Santa Cruz | sc-8654 | 1:2000 (WB) |

*IF: immunofluorescence; WB: Western blotting
